# Trait-based mechanistic approach highlights global patterns and losses of herbivore biomass functional diversity

**DOI:** 10.1101/2021.10.19.464976

**Authors:** Fabio Berzaghi, Dan Zhu, John Alroy, Philippe Ciais

## Abstract

Mammalian herbivores strongly shape Earth system processes. A lack of global dynamic models of mammal populations limits our understanding of their ecological role at large scales and the consequences of their extinctions. We developed a mechanistic global model of terrestrial herbivore populations simulated with 37 functional groups defined by analyzing eco-physiological traits of all extant herbivores (n = 2599). We coupled this model with a global vegetation model to predict herbivores’ biomass maximum potential (pre-industrial) and at present. Natural ecosystems could have sustained a wild herbivore wet biomass of 330 Mt, comprised of 192 Mt by large (body mass > 5 kg) and 138 Mt by small species. Anthropogenic activity reduced large herbivores biomass by 57%, estimated now at 82 Mt; consequently, small herbivores now dominate global herbivore biomass with 98 Mt (-29%). Losses vary greatly across climatic zones and functional groups, suggesting that size is not the only discriminant feature of biomass decline. Actual evapotranspiration is the most important driver of total, large, and small herbivore biomass and explains 64%, 59%, and 49% of its variation, respectively. Energy and water dependency is high for large herbivores, whose biomass is greater in hot and wet areas, challenging the notion that large herbivore biomass peaks in savannas only. Outside Africa and the Tropics, potential-biomass hotspots occur in areas today dominated by humans; this could undermine the recovery of larger species in certain areas. These herbivore biomass estimates provide a quantitative benchmark for setting conservation and rewilding goals at large spatial scales. The herbivore model and functional classification create new opportunities to integrate mammals into Earth System science and models.

## Introduction

Terrestrial mammalian herbivore species shape ecosystems, alter vegetation and soil functioning, change plant biodiversity, and accelerate carbon and nutrient turnovers (Malhi et al., 2016; Schmitz et al., 2023). In particular, herbivores heavier than 5 kg (Berzaghi et al., 2018) (also called large) influence carbon cycling and biosphere-atmosphere feedbacks (Berzaghi et al., 2023; Norris et al., 2020; Schmitz et al., 2023). Much less is known about the role of lighter terrestrial herbivorous mammals (body mass < 5 kg), in spite of their effects on plant diversity, vegetation structure, and soil physio-chemical properties (Bomske & Ahlers, 2021; Gordon & Prins, 2019; Poe et al., 2019). Light mammals (also called small) likely perform very different functional roles than larger mammals (e.g., burrowing). Range contractions and local extinctions prompt for a better understanding of mammals’ multifaceted ecological role and repercussion on ecosystem functioning. This knowledge is needed to guide conservation and trophic rewilding and to restore mammals-driven ecosystem services (e.g., reducing wildfires, increasing nutrients cycling, and enhancing carbon storage) which could contribute to mitigating climate change (Cromsigt Joris P. G. M. et al., 2018; Malhi et al., 2022; Schmitz et al., 2023). Extinction or decline is evaluated often in terms of changes in species richness, absolute abundance, or density per unit area within a locality. However, quantifying mammalian herbivore biomass across the whole spectrum of body mass is critical for assessing mammals’ contribution to past and present ecosystems and Earth System processes. Specifically, estimates of functional biomass, which is animal biomass classified in functionally-diverse groups, are lacking but are greatly needed to determine the intensity of animal-environment interactions at various eco-spatial scales (Berzaghi et al., 2018; Gaston et al., 2018; Norris et al., 2020). Trait-based theory and empirical methods to calculate the functional biomass of mammals have been applied to a few groups: African savanna large grazers (Pachzelt et al., 2015) and sub-Saharan medium-large ungulates (Harfoot et al., 2014; Hempson et al., 2015). By comparison, plant and marine ecologists have developed global trait-based classifications to mechanistically simulate various organisms (e.g., plants, plankton, fish) dynamics, biomass, and contribution to biogeochemical cycling (Aumont et al., 2018; Krinner et al., 2005; Pinti et al., 2023). Trait-based classifications obviate the need to independently model the ecophysiology of thousands of species. Instead, well-structured groups are manageable for implementation in large-scale terrestrial biosphere models. These methodologies are now central in macroecology, Earth System science, and environmental policy (Aumont et al., 2018; Berzaghi et al., 2020; Rosenzweig et al., 2017). Global mechanistic models of mammalian herbivores would add herbivory to global Earth System models, elucidate the effects of past mammal extinctions, and inform future conservation strategies (Norris et al., 2020).

Here we present a novel methodology for integrating mammalian herbivores in Earth System studies and for estimating herbivore biomass across size and functional classes. We develop the first global classification of extant mammalian herbivore species into tractable herbivore functional types (HFTs) (Hempson et al., 2015; Pachzelt et al., 2015). The HFT classification is defined by four traits: diet type, adult body mass, digestive system (hindgut or foregut), and folivory type (grazers, browsers, and mixed-feeders). We implemented this new HFT classification in an eco-physiological mechanistic model coupled with the output of ORCHIDEE, a widely-used terrestrial biosphere model (Krinner et al., 2005). The herbivore model, hereafter REMAP (REproduce MAmmal Populations), simulates the carrying capacity of all herbivores, grouped in HFTs, as a result of reproduction, metabolism, growth, mortality, and competition for food. We used REMAP to simulate present-day potential-maximum biomass distribution under natural conditions: pre-industrial climate, natural habitat only, and no widespread human disturbances such as hunting. Lastly, we used these results to quantify changes in functional biomass caused by habitat and range loss, pinpoint functional groups and regions in greater need for protection, and investigate the main environmental drivers influencing herbivore biomass productivity.

## Methods

### Methodology overview

Large-scale mechanistic models of highly biodiverse organisms are computationally intensive and require heavy parameterizations. Consequently, global models of plants and animals do not simulate eco-physiological processes of thousands of species, but instead they use functional groups to simulate species with similar traits (Aumont et al., 2018; Harfoot et al., 2014; Krinner et al., 2005; Pinti et al., 2023). To simulate the global population dynamics of extant mammalian herbivores, we first developed a classification of herbivore functional types using clustering and statistical analysis applied on mammals’ trait data gathered from different sources. These HFTs were then incorporated in a mechanistic model that simulated the global distribution of pre-industrial (only natural habitat and no intensive hunting) biomass of each HFT as a function of environmental conditions and daily changes in plant biomass availability. An additional simulation produced biomass estimates accounting for present-day range and habitat loss. REMAP biomass emerge from the HFTs population dynamics, including mortality, and the availability of and competition for food among HFTs. Eco-physiological processes in REMAP are described through mathematical equations based on HFT-specific parameters, most of which are calculated from body mass scaling equations (Pachzelt et al., 2015; Zhu et al., 2018). The model results were validated against empirical data of mammal biomass compiled from different sources (*Data* section). An additional validation was performed by complementing empirical data with estimates from regression analysis (Santini, Isaac, Maiorano, et al., 2018); this was needed for smaller species which were data deficient.

### Data section and analysis

#### Mammals’ trait data

We gathered extant terrestrial mammalian herbivore species traits data from available datasets: Phylacine (n = 5831), Mammal Diet 2 (n = 6625), Elton Traits (n = 5400), and PanTHERIA (n = 5416) (Faurby et al., 2018; Gainsbury et al., 2018; Jones et al., 2009; Wilman et al., 2014). These datasets do not include semi-domesticated or range species, which were not part of our study. We harmonized taxonomic differences between data sets by using the Phylacine standardized synonym table. These datasets include life-history traits (body mass, diet preferences, longevity, etc.) for all known mammal species.

#### Empirical mammal population density & biomass

We compiled a dataset of mammal density estimates from two main sources (Hatton et al., 2015; Santini, Isaac, & Ficetola, 2018) that we supplemented with 433 records we found in literature for a total of 7713 records (complete list of density data in Appendix). This dataset included 440 species, which represent roughly 50%, 15%, and 7% of the species that we later define as, respectively, large (n = 238), small (n = 108), and micro (n = 94). The mammal density data were collected between 1950-2020 in locations such as national parks or protected areas where human disturbance should be less than outside protected areas (Santini, Isaac, & Ficetola, 2018). The data had geographic and taxonomic gaps, small and micro mammals were particularly underrepresented, because only a fraction of all species present in each location were censused (Santini, Isaac, & Ficetola, 2018) (Fig. S1). This lack of taxonomic coverage reduces considerably the estimates of total animal biomass per km² (Fig. S1). Various statistical methods have been used to estimate mammals density/biomass to complement empirical data (Greenspoon et al., 2023; Santini, Isaac, Maiorano, et al., 2018; Vidal-Cordasco et al., 2022). We thus generated density estimates for the missing locations and species using a spatial linear mixed-effects regression following (Santini, Isaac, Maiorano, et al., 2018). The linear mixed-effects regression was built using the following predictors: body mass, grass and tree NPP from MODIS (Heinsch et al., 2003), precipitation of the warmest quarter from WorldClim2 (Fick & Hijmans, 2017), and random effects to account for taxonomy and spatial pseudo-replication. The linear mixed regression captures some of the influence of bioclimatic factors on intra & inter-species density variability (Santini, Isaac, Maiorano, et al., 2018). REMAP generates community-wide biomass estimates; thus, the necessity to complement the empirical data with independently-estimated data. We only estimated density in grid cells where species are present in accordance with the *Phylacine* “present natural” distribution maps (Faurby et al., 2018). “Present natural” maps are constructed from IUCN maps and represent estimates of presence if there were no anthropogenic disturbances or pressures and do not account for invasive or introduced species. These natural conditions matched our model simulation conditions (*REMAP* section). The empirical data and the mixed-effects model estimates were combined to derive biomass per km² by multiplying density by 80% of the body mass reported in Phylacine to account for sexual dimorphism and juveniles (Fairbairn et al., 2007).

#### Biomass loss estimates

We estimated the amount of biomass lost due to geographic range reduction and habitat loss by following a two-step procedure. We first simulated with REMAP the maximum potential herbivore biomass according to the “current” range maps provided by the *Phylacine* database (Faurby et al., 2018) (see *REMAP gridded inputs* section for more details on the setup of the simulation). These maps are largely based on the species ranges estimated by IUCN in 2016. However, these maps alone are not suitable to estimate area-based loss of biomass because they do not account for the fraction of the landscape which is not natural habitat. We thus estimated the fraction of habitat loss by aggregating at 2° resolution the artificial habitat layer of a global map of terrestrial habitats (Jung et al., 2020). In the original map the artificial habitat included urban areas, plantations, and agricultural lands but not rangelands because they are not present in the IUCN habitat classification scheme. Thus, we might underestimate the artificial habitat in each cell and we acknowledge that at times wild herbivores use crops as food source. We then used the artificial habitat map to calculate the fraction of remaining biomass in each grid cell after habitat loss. Hunting likely further reduces biomass, particularly of large species in tropical ecosystems (Benítez-López et al., 2019). We found no global gridded data allowing to estimate biomass loss due to hunting, but some should be accounted by the range and habitat loss calculations.

#### Correlation analysis between climate data and biomass

Yearly averages of plant evapotranspiration, precipitation, temperature, and precipitation seasonality were obtained from TerraClimate and WorldClim 2.1 (Abatzoglou et al., 2018; Fick & Hijmans, 2017) (further details in Appendix S1). Climate data were regressed against modeled biomass with linear and second order polynomial models.

### Global classification of mammalian herbivores

We used Phylacine as our taxonomic and diet reference database, as it is the most recent database and uses phylogeny to better estimate species diet composition (Faurby et al., 2018). We selected all terrestrial mammalian species whose diet is composed of at least 80% plant material (n=2599, Appendix S1 for more information on excluded species). For each species we determined diet composition in terms of the percentage of leaves, fruit, and seeds consumed according to published data sets of mammals diet (Kissling et al., 2018; Wilman et al., 2014). Fruit, seeds, and leaves were chosen because they cover the vast majority of mammalian herbivores’ diet and are most frequently simulated in vegetation models (further details on methodology in Appendix S1). We also determined digestive system (hindgut or foregut) and, only for species consuming leaves, a folivory type: grazer, browser, or mixed-feeder. We defined “grazer” as consumers of primarily herbaceous plants (“grass” hereafter), “browsers” as consumers of woody plants (“browse” hereafter), and “mixed-feeders” as consumers of both (further details on methodology in Appendix S1).

The HFTs were created following a nested four-step approach in which each species was assigned to a functional type based on its dietary preferences, body mass, folivory type (only for leaf eating species), and digestive system (Fig. S2).

Step 1: we assigned species to dietary macro-groups by applying hierarchical cluster analysis (R package “factoextra”) using the three plant organs as data input. We used the “agnes” algorithm as it provided the tightest clusters that were evaluated using silhouette width (average distance between clusters), Dunn index, and standard deviation. We found eight to be the optimal number of clusters (dietary macro-groups) based on statistical evidence and ecologically-meaningful diet types. These groups were: Leaf, Leaf & fruit, Fruit & leaf, Fruit, Leaf & seed, Seed & fruit, Seed, and Generalist (Fig. 1). These groups represent folivores (Leaf), granivores (Seed), and frugivores (Fruit), and all of the combinations highlighted by the analysis (Fig. S3). Less than 3% of all species were at the edges of each cluster (negative or low silhouette width). These species were reassigned to the group with the most similar dietary preference. This process improved the clusters compactness by increasing the silhouette coefficient and reducing the standard deviation of diet components.

**Fig. 1.**
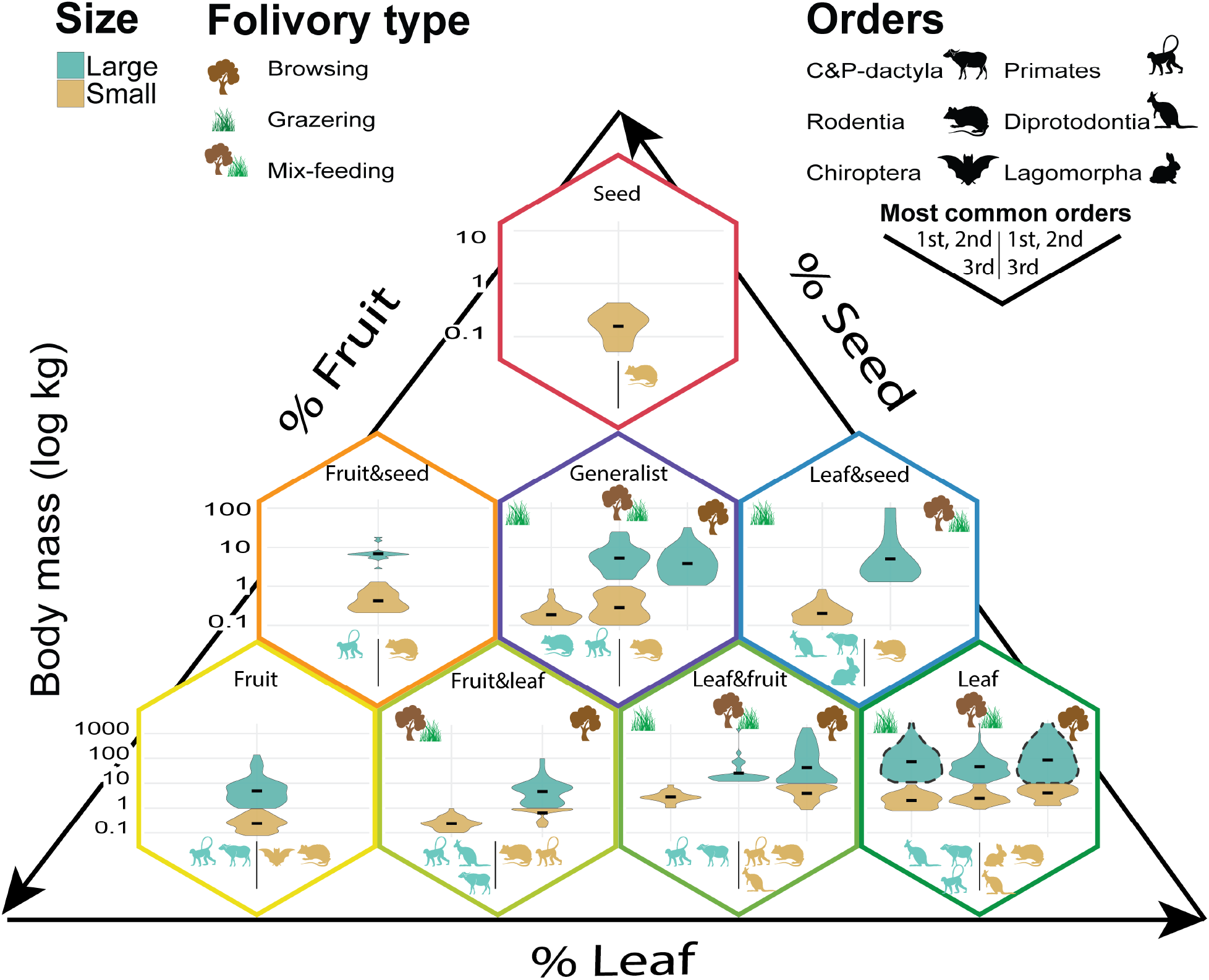
Characteristics of HFTs within the ternary space Leaf-Fruit-Seed. Each violin plot corresponds to one HFT. Animal icons show the 1-3 most common orders among large and small HFTs within each diet macro-group based on number of species. C&P stands for Cetartiodactyla and Perissodactyla. Foregut fermenters are indicated with a dashed outline, the other HFTs are hindgut fermenters. Micro HTFs, not shown in this graph, have the same traits of small HFTs aside from body mass and are mostly rodents (83%) and bats (10%).

Step 2: Within each of the eight diet macro-groups we assigned species to three size classes: micro (1398 species, mean 0.13 kg, range 0.004-0.05 kg), small (726 species, mean 1.6 kg, range 0.05-10 kg) and large (475 species, mean 92.4 kg, range 1-3520 kg) animals, only the Seed group did not contain large species. Division in size classes was necessary because body mass in mammals spans various orders of magnitude and underlies significant eco-physiological differences such as longevity, litter size, and litter per year. Change-point analysis was used to determine significant differences in mean body mass among species in each diet group. Change-point analysis is a technique used to detect trend changes in timeseries but when applied to ordered data (Beaulieu & Killick, 2018) it helps identify groups having similar mean values. Along with the results from the change-point analysis, we also considered biologically-relevant information, such as taxonomy, to minimize arbitrary species assignments. A size threshold was determined within each macro group (Fig. S4. Consequently, the thresholds vary in each diet group according to the species present in it. As a result, thresholds can slightly overlap between size groups across diet groups. Previous studies have proposed slightly different body mass categories, but did not have the fine-grained functional trait resolution that we propose here (Benítez-López et al., 2019; Vidal-Cordasco et al., 2022).

Step 3: this step was only necessary for species that are partly-or exclusive leaf consumers; thus, it applied to all dietary macro-groups excluding Seed, Fruit, Fruit & seed. Species within each diet/size were grouped according to their feeding preference for herbaceous, woody, or mixed vegetation.

Step 4: the final nested grouping was based on digestive system, either hindgut or foregut, and was applied to all diet/size/folivory type groups. Non-ruminant foregut fermenters were treated as foregut fermenters because fermentation takes place in the same part of the stomach and specific model parameters for non-ruminants were not available (Shipley et al., 1999).

The key traits used in step 3 and 4 traits influence, among other features, maximum daily intake (eq. 1 and 2 shown later) and type of plant consumed (grasses, trees, or both). After step 4, each species was assigned to an HFT (list of species in Data S1) defined by: 1) diet composition (percentages of leaf, fruit, and seed in diet) determined by the average of all species within each HFT; 2) size class (large, small, and micro); 3) folivory type (only for exclusive or partly-folivores); 4) digestive system (hindgut or foregut). The most common trait combinations, in terms of number of species and geographic range, were retained to avoid creating hundreds of HFTs. Because more than 85% of species were hindgut fermenters, folivory type was used to reduce the number of HFTs by reassigning species to the most functionally-similar HFT. Consequently, some species were reassigned to HFTs with a different folivory type, such as placing browsers or grazers into a mixed-feeder HFT (List of species and respective HFT in Appendix). In total we had 37 HFTs: 11, 13, and 13 for large, small, and micro herbivores respectively (see Table S1A for small and large HFTs). The HFTs for micro mammals differed from small HFTs only in body mass; whereas diet composition, digestive system, and folivory type were the same. In REMAP simulations, micro mammals’ biomass was only 5% of the total biomass. Thus, for result presentation purposes, their simulated biomass was added to the biomass of small HFTs.

### REMAP: a mechanistic model of herbivores ecophysiology

The above-described HFTs were implemented in REMAP, a mechanistic model that simulates ecological, physiological, and demographic processes of different HFTs competing for food. These processes are simulated with mathematical equations that allow to calculate each HFT maximum energy intake and energy expenditure based on physiology, body mass allometric scaling, and air temperature (Zhu et al., 2018). Body mass of each HFT varies spatially according the geographic range of its species. Body mass is calculated per HFT per grid cell as the average of all species within the cell weighted by total number of individuals (Zhu et al., 2018). This avoids body mass allometric scaling calculations biased towards larger species in HFTs containing large variations in body mass (see Appendix S1). It also reduces the need to have too many size classes which become computationally burdensome and increase the number of parametrizations. Herbivore functional types consume the available plant biomass (leaves, fruits, seeds) simulated by the vegetation model ORCHIDEE (described below) and convert the energy surplus into fat storage. HFTs compete for food and prioritize feeding on fresh biomass otherwise they resort to eating litter or starve if no food is available. Fat storage determines annual birth rates and influences mortality rates because fat is converted to energy during periods of starvation. Conceptually, fat storage is used as a proxy for energy stored in tissue that is mobilized to perform different functions. In reality, other tissues such as muscles and bones are mobilized during starvation and reproduction but fat is the primary one (Devlin, 2011; Heldstab et al., 2017). Predators are not explicitly modeled but their effect on increased herbivore mortality is captured by using a carrying capacity parameter in a logistic equation. Non-consumptive effects of predators on herbivores, such as limiting access to resources (i.e., landscape of fear) are not included. Even though most equations are based on Zhu et al. 2018, most parameters were re-estimated from trait data to accommodate for the wide variation in eco-physiology within the new HFTs (Tables S1A, S2-S3). The key estimated parameters include: maximum fat storage, aging mortality, establishment rate, daily intake and food energy content, carrying capacity, and energy expenditure (Table S2). Compared to Zhu et al. 2018, competition for food among HFTs and flexible diets are key added features along a new establishment scheme to account for small species breeding frequently throughout the year. A more detailed description of these processes and their equations is provided in the Appendix S1.

#### Model setup and inputs

REMAP spatial resolution is determined by the resolution of its inputs, notably: a) gridded time series of production and turnover rate of plant organs (leaf, fruit, and seed) and b) HFT body mass maps. a) Time series of plant organs biomass can be derived from remote-sensing products or from the output of global vegetation models. For our simulations we generated a 300-year time series, enough to reach a steady state, of plant organs at 2° resolution using the ORCHIDEE (Organizing Carbon and Hydrology In Dynamic Ecosystems) vegetation model (Krinner et al., 2005). ORCHIDEE is a widely-used dynamic global vegetation model and provides net primary productivity (NPP) allocation and turnover rate at a daily time step containing fresh leaves and litter from herbaceous (grasses) and woody (trees) vegetation, fruit, and seeds (see Appendix S1 and Krinner et al. 2005 for full description of ORCHIDEE). Even though ORCHIDEE and REMAP are not fully coupled, the available plant biomass pools are updated daily to account for animal consumption. A future application to utilize the full potential of REMAP could include a fully-coupled simulation in which the feedbacks between animals and plants would allow a more precise estimation of the available biomass pools and nutrient soil availability. For example, the removal of leaf biomass would change water use and carbon allocation of plants and defecation would affect NPP through nutrient distribution. B) Maps of HFT average body mass are a prescribed input in REMAP. We generated two global maps of HFT body mass at 2° spatial resolution to match the spatial resolution of ORCHIDEE. The gridded body mass maps were created according to the “present natural” and the “current” distribution maps previously described in the data section (Fig. S5). The “present natural” body mass maps were used to simulate potential herbivore biomass for pre-industrial natural conditions. Whereas, the “current” body mass maps were used to simulate herbivore biomass accounting for species range contraction.

REMAP output consists of time series of population densities (individuals/km²) for each HFT. Densities were averaged over the last 50 years of the simulation because populations can fluctuate over time due to demographic and ecological processes. Densities were then multiplied by HFT body mass to obtain live biomass wet weight (kg/km²) for each HFT at 2° degrees resolution.

### REMAP validation data

We validated REMAP “current range” predictions for large HFTs against field-based densities for locations (n = 81) where at least 70% of all large species were censused, providing an acceptable estimation of total large herbivore community biomass and geographic coverage. We also compared REMAP with empirical densities complemented with estimates generated with the regression analysis described in the data section. We calculated the mean average error (difference between modeled and empirical biomass), Pearson correlation, and computed the residuals by subtracting empirical biomass from the REMAP-modeled biomass.

## Results

### Global mammalian herbivores classification

We classified all extant terrestrial mammalian herbivores (2599 species) into eight (Fig. 1 and S3). These eight groups showed minimal cluster overlap (Fig. S3A) and no diet overlap in terms of the standard deviation of their diet components (Fig. S3B). Taxonomic diversity and body mass varied greatly among diet groups (Figs. 1 and S4, and Table S1A). From the eight dietary macro-groups, HFTs were derived to reflect empirically-observed relationships between adult body mass, digestive system, and folivory type (Fig. 1 and Table S1). The HFTs are discrete groups but their parameters in REMAP cover a continuous trait space because parameter values are determined with continuous functions and diet is flexible within the limits of each HFT (Fig. S3B, Methods). These broadly-studied traits govern most mammals’ eco-physiological processes but had not been used jointly to create distinct trait profiles that are also compatible with terrestrial biosphere models. There were three size classes within each macro-group: large, small, and micro. Large and small body mass overlap slightly because size thresholds were determined independently for each diet macro-group (Fig. S4 and Methods). Hindgut fermenters (n=2245) and foregut fermenters (n = 354) were separated. Foregut fermenters are generally heavier than hindgut fermenters (Fig. 1). Among obligate-leaf consumers, grazing is most common (45% of species) followed by mixed-feeding (30%) and browsing (25%). The separation of species by these traits led to the definition of 37 HFTs (Fig. 1), including 11 HFTs representing large herbivores and 13 HFTs for small ones (see note on micro HFTs in Fig. 1 legend and Methods). Even though we separated HFTs by size classes, intra-HFT body mass was spatially variable as it reflects geographic ranges (Fig. S5). This spatial variability was accounted for in our simulation of mammalian biomass (Methods)

### Global patterns of potential herbivore biomass

The total global REMAP-modeled biomass of all herbivores was ∼330 Mt (live weight), of which ∼192 Mt comprised large HFTs and ∼138 Mt small HFTs (Fig. 2A). Micro HFT biomass was ∼5% of the total biomass. These figures represent the potential carrying capacity of present-day herbivores without habitat loss or other anthropogenic pressures. Total modeled biomass roughly halved between warmer and wetter climates and between drier and colder ones (Fig. 2B). Most biomass (81%) consisted of Leaf and Leaf & fruit consumers (Fig. 2B). To our knowledge, directly comparable results do not exist. Madingley, a mechanistic general ecosystem model, produced total mammal biomass estimates of the same order of magnitude but with no differentiation between size classes and with markedly different spatial patterns (Enquist et al., 2020). This is not surprising, as REMAP and Madingley use different approaches. Madingley models generic endotherm herbivores without specifying diet or digestive system, but uses a cohort system (Harfoot et al., 2014). In Madingley, food availability is based on the internally-calculated autotroph biomass. Instead, food availability is an independent input from a dedicated vegetation model (i.e., ORCHIDEE). Incorporating herbivore traits such as digestive system, diet, and geographic distribution of actual species are fundamental to predict herbivore biomass and will be useful to assess megafauna extinctions as extinct species had distinctive traits (Pringle, 2020). The modeled biomass was within similar ranges to the 380 Mt (270 Mt for large and 110 Mt small HFTs) estimated independently with gap-filled empirical data (Fig. S6). Large HFTs biomass was also in reasonable agreement with observed biomass in locations where enough species were censused (Fig. 3). The agreement between modeled and empirical biomass was highest in high-biomass regions: tropical, temperate, and cold (Figs. 2, S6; climate zones follow the Koppen-Geiger classification in Fig. S7A). Biomass was underestimated mostly in some high latitude regions and in or near large deserts; however, residuals in most continents were normally distributed around zero or -0.5 (Fig. S6). In most of these areas, food availability in the REMAP input was low (Fig. S8A) and the empirical data were sparse or absent (Fig. S1 and Methods) making punctual comparisons difficult. For example, empirical biomass estimates of two hyrax species equate to 191 Mt alone when extrapolated across their ranges. This represents 63% of empirically-predicted small HFT biomass and results in high negative residuals in Africa arid areas (Fig. S6A). Other discrepancies are expected between empirical and REMAP data as the former are often a snapshot in time whereas the latter are an average of population dynamics through time.

**Fig. 2.**
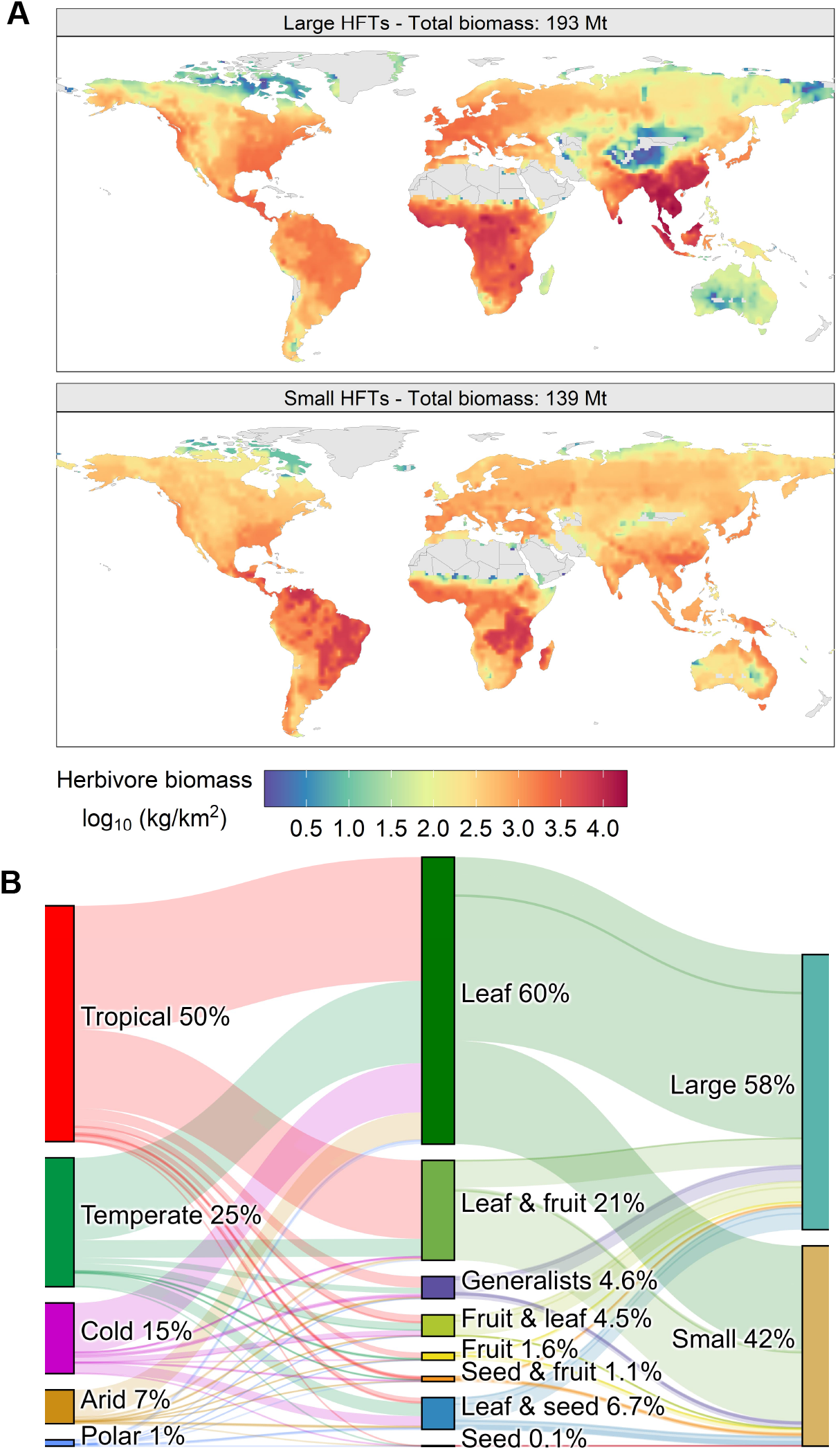
Distribution of present-day modeled biomass. (A) Spatial distribution of modeled biomass under natural conditions (no artificial habitats and intensive hunting) for the two main mammal size classes; body mass of large (> 1-10 kg) and small (< 1 kg) varies spatially (Fig. S5). (B) Relative biomass distribution across climate zones and size classes.

**Fig. 3.**
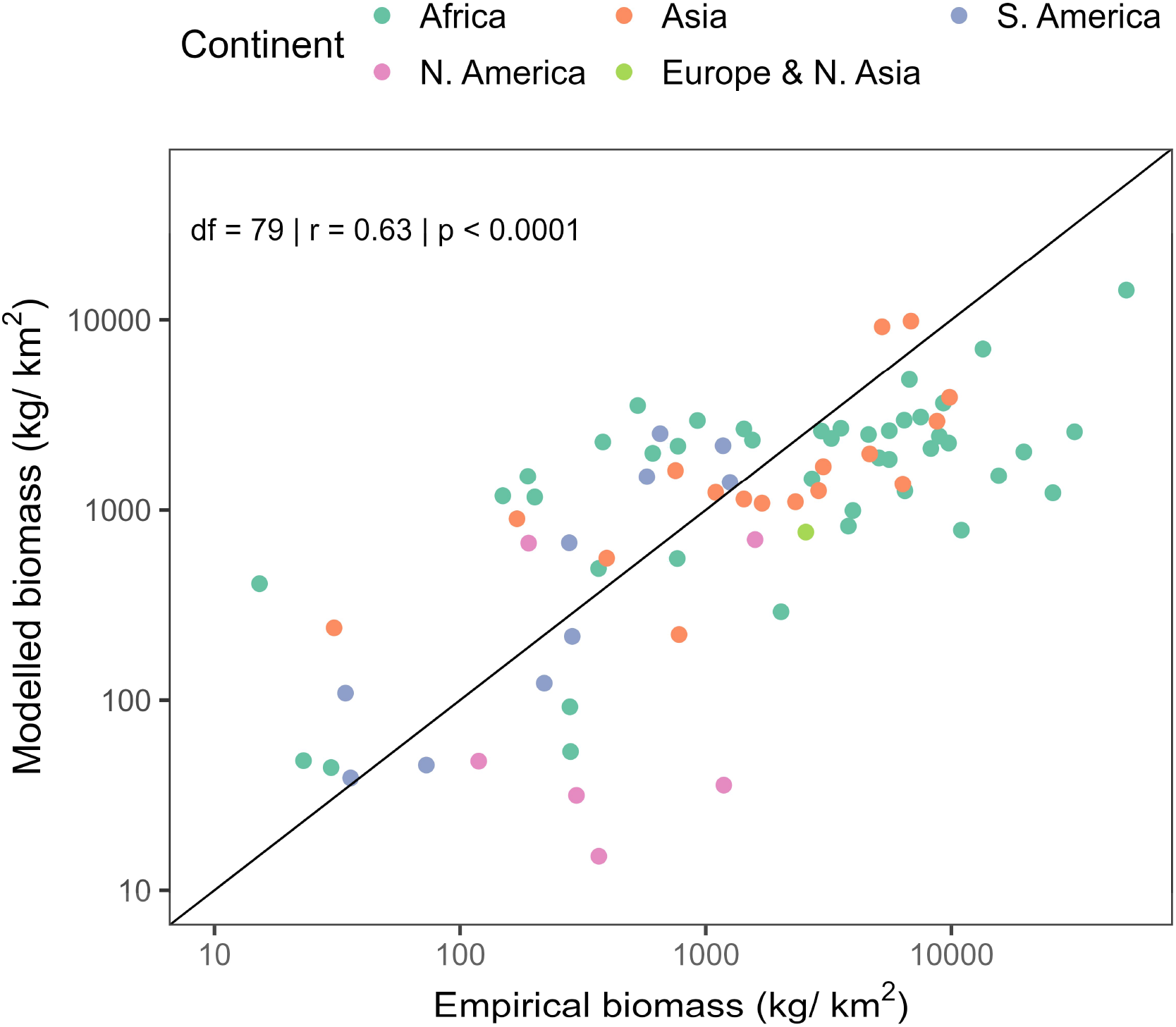
Comparison between modeled large HFTs biomass and empirical biomass of large herbivores. Modeled biomass refers to “current range” simulations accounting for anthropogenic pressures (range reduction and habitat loss); empirical biomass is only based on field measurements in locations where more than 70% of species were censused (Methods). Biomass was log transformed, the dashed line represents the 1–1 line, and pearson correlation test (two sided) was used to calculate *r* and *p*.

Notable differences in spatial patterns between class sizes were found in the northern cold climates, tropical, and sub-tropical areas (Fig. 2A). For instance, in polar climates, small herbivore biomass was four times higher compared to large herbivores and the two groups had similar total biomass in cold and arid climates (Fig. 2B). In terms of biomass distribution across functional types, small and large Leaf consumers were the most widely distributed and most variable in biomass per area (1-10,000 kg/km², Figs. 4 and S9). Leaf consumers biomass peaked in southeast Asia (5000-10,000 kg/km²) due to high abundance of large mixed-feeders, which include the Asian elephant (*Elephas maximus*) (Fig. 4 and S9A). Generalists were widespread but with much lower biomass (10-500 kg/km²) peaking in sub/tropical areas and in the southeastern USA (Fig. 4). Large Fruit & leaf and Leaf & seed were the only groups whose biomass was higher (50-500 kg/km²) in temperate and cold areas than the tropics (Figs. 4 and S9A). These groups include the American black bear (*Ursus americanus*) and the wild boar (*Sus scrofa*), which are the most widespread species in North America and Europe.

**Fig. 4.**
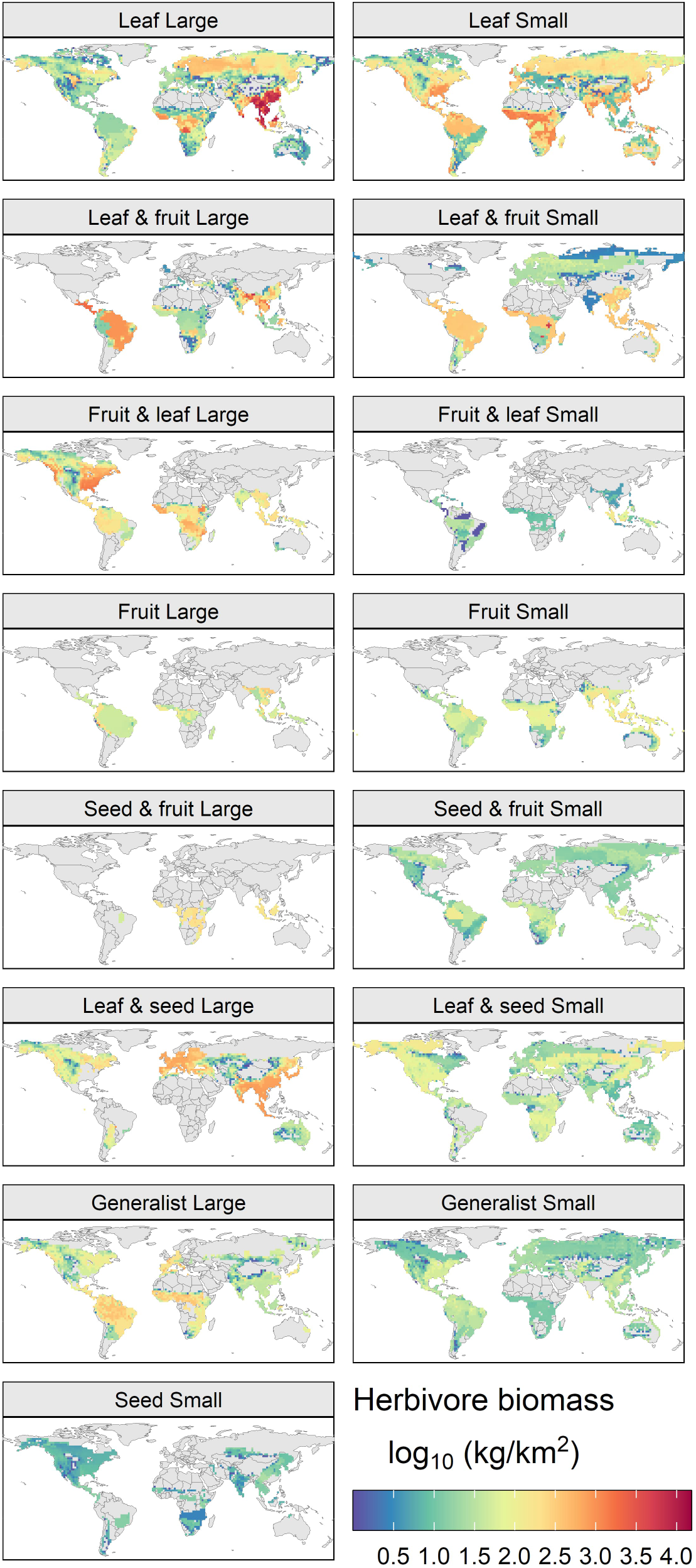
Modeled biomass of dietary macro-groups under the natural conditions scenario. The biomass of large (left column) and small (right column) HFTs within the same macro-group was aggregated. The Seed micro-group exists only for small HFTs.

### Global drivers of herbivore biomass

Small and large herbivores biogeographic patterns were dissimilar (Fig. 2A). Large herbivore biomass was higher in warmer and wetter locations associated with high plant actual evapotranspiration (R² = 0.59, p < 0.001), precipitation (R² = 0.53, p < 0.001), and temperature (R² = 0.30, p < 0.001) (Fig. S10A-C). Instead, small herbivore biomass was more evenly distributed and less correlated to evapotranspiration (R² = 0.49, p < 0.001), precipitation (R² = 0.41, p < 0.001), and temperature (R² = 0.15, p < 0.001) compared the biomass of large herbivores (S10A-C). No apparent correlation was found with precipitation seasonality (Fig. S10D). Evapotranspiration explained even more variability (R² = 0.64, p < 0.001) when regressed against total biomass, small and large biomass combined, whereas the explanatory power of the other climatic variables did not increase (Fig. S11). However, correlating biomass modeled in “current conditions” (accounting for anthropogenic disturbances) with evapotranspiration and precipitation resulted in a lower fit (Fig. 5; R² = 0. 49 and R² = 0. 50, respectively). Biomass estimates from observations seemed to also suggest linear log-log relationship between biomass and evapotranspiration (Fig. 5A, R² = 0. 20, p < 0.001) and a more left-skewed quadratic relationship with precipitation (Fig. 5 B, R² = 0. 26, p < 0.001), but more data would be needed to better assess this empirically. Previous analyses showed that primary productivity and precipitation seasonality influence the density of mammals at the species level (Santini, Isaac, Maiorano, et al., 2018). However, here we evaluate biomass at the community level as a result of trophic interactions and direct comparisons with incomplete density surveys are challenging. These previously-described biogeographic patterns and the evapotranspiration-biomass correlations suggest that, globally, large herbivores are more energy and water dependent than smaller ones, many of whom can extract water from plants through selective feeding (Gordon & Prins, 2019). These conclusions are compatible with previous observations of water-size class correlations in African herbivores (Hempson et al., 2015). Our findings, along with other recent articles (Greenspoon et al., 2023; Vidal-Cordasco et al., 2022), showed that herbivore biomass could reach considerable amounts in certain tropical and wet areas. This challenges the assumption that large herbivore biomass peaks in savannas and at intermediate moisture levels. In the Congo Basin, forest elephant densities have been reported at 3.5 km² or ∼10,000 kg/km² (Poulsen et al., 2017), to which we should add ungulates, apes, and hippos, which are semi-aquatic but feed primarily on land.

**Fig. 5.**
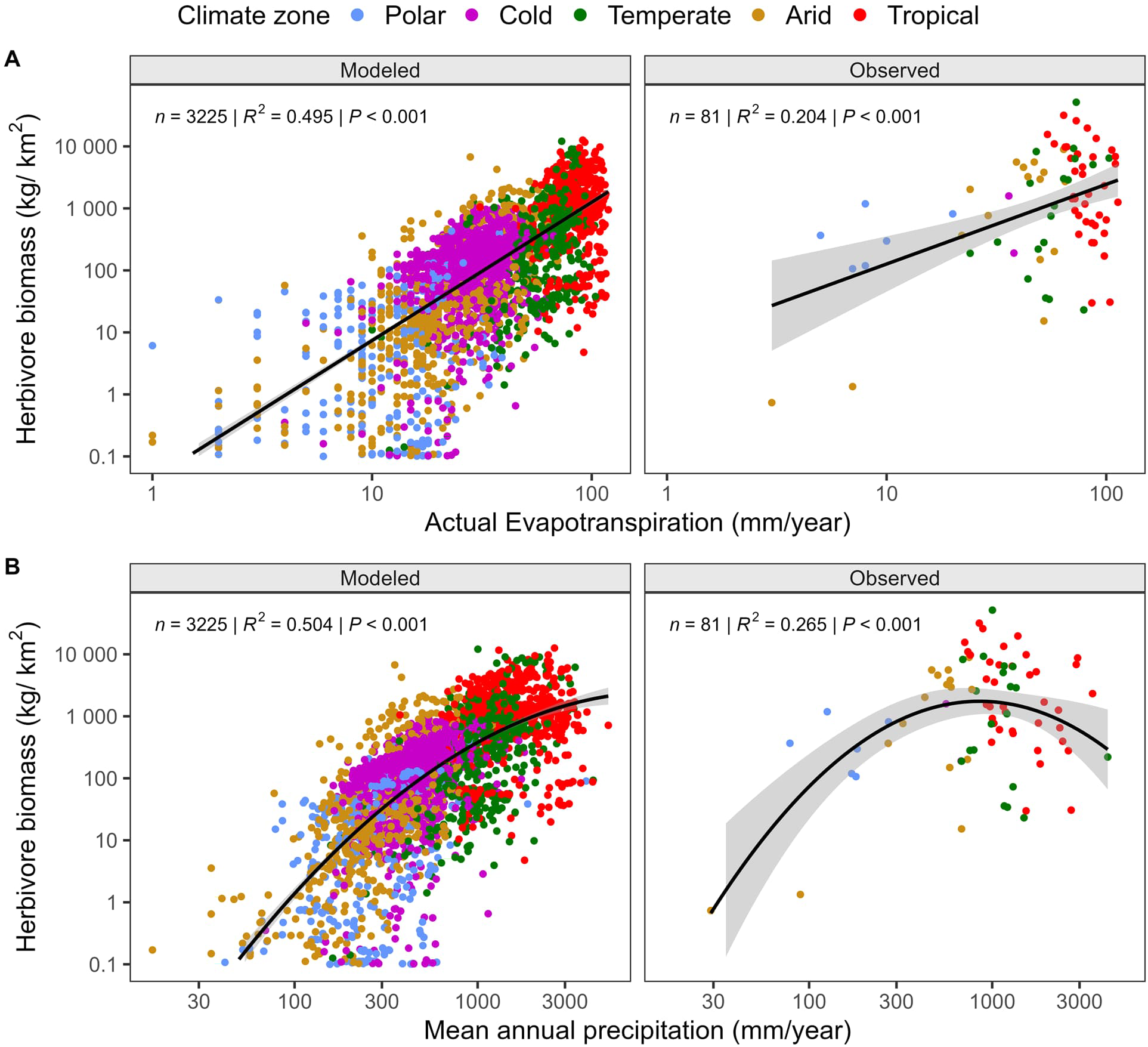
Correlations between modeled or observed herbivore biomass and climatic factors. Modeled biomass refers to the “current range” simulations accounting for anthropogenic pressures (range reduction and habitat loss); empirical biomass is only based on field measurements in locations where more than 70% of species were censused (Methods). Biomass, (**A**) Actual evapotranspiration, and (**B**) mean annual precipitation were log transformed.

Furthermore, the weaker correlation between herbivore biomass and evapotranspiration when accounting for anthropogenic pressures (Fig. 5, S10, and S11) suggests that: 1) using current population estimates of mammals’ density or biomass might lead to weaker correlations with environmental variables because many populations are not at their potential densities; 2) evapotranspiration could be used to predict potential maximum biomass in non-sampled areas or for choosing locations more suitable for rewilding.

### Consequences of range and habitat loss

By modeling present-day biomass accounting for mammals’ range reduction and habitat loss, we estimated that only 82 Mt of large and 98 Mt of small herbivore biomass remain. These numbers suggest a potential loss of almost ⅔ (-110 Mt) of large herbivore biomass and ⅓ (-40 Mt) of small herbivores (Fig. 6). Observation-based estimates for 238 large herbivores species provide a total biomass of ∼25 Mt, but their spatial coverage is very fragmented (Fig. S1), suggesting a large underestimation. Both our results, from model and observations, suggest that recent estimates of ∼22 Mt accounting for all terrestrial mammals across their current ranges, might also underestimate mammals’ biomass (Greenspoon et al., 2023). Absolute biomass losses across climate zones, functional groups, and class sizes are similarly proportioned to the distribution of their total potential biomass with tropical and temperate Leaf and Leaf & fruit groups showing the largest losses (Fig. 2B and Fig. 6C). However, the analysis of relative losses reveals that even if absolute biomass losses were highest in tropical and temperate zones, relative losses are consistently around 25% or more for most functional groups and in all climate zones (Fig. 6B). Certain functional groups, in particular Fruit and Leaf & seed, have lost around 50% or more of their biomass (Fig. 6B). Additionally, in all but polar climate zones, small and large herbivores biomass have seen similar relative losses. This suggests that functional biomass has been greatly diminished not only for large herbivores in tropical and temperate areas, but evenly across the board.

**Fig. 6.**
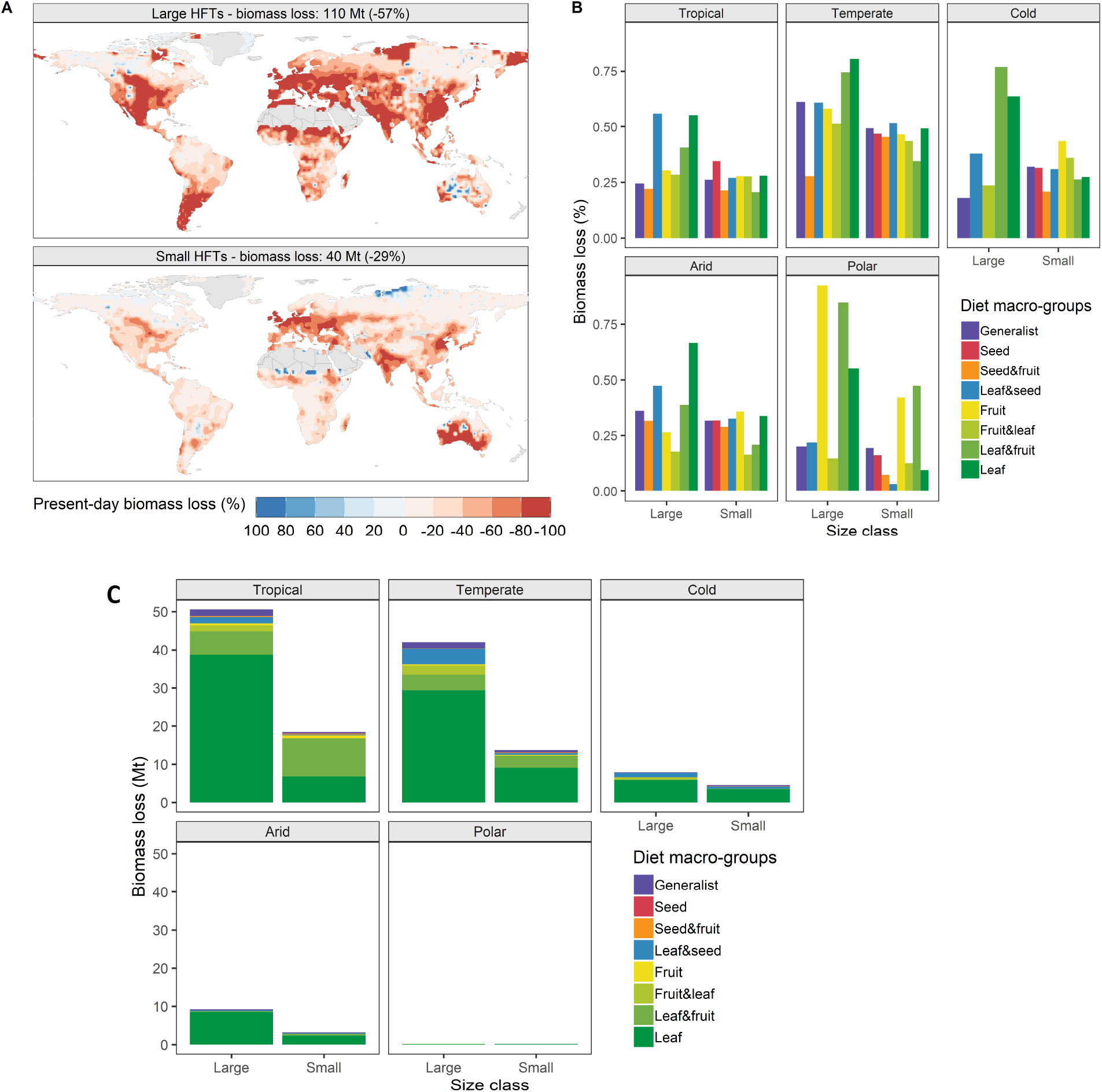
Absolute and relative losses in herbivore biomass due to range contractions and habitat loss. **A** Percentage change in herbivore biomass calculated from the difference between the potential biomass in natural conditions (no intensive anthropogenic pressure) and “current range” conditions. **B** Relative biomass loss by climate zone, class size, and functional group. **C** Absolute biomass loss in Mt of live weight.

Outside of Africa, predicted hotspots of potential large herbivore biomass coincided with human-occupied landscapes: western Europe, southeastern China, central North America, and India (Figs. 2A, 4, and S7C). These areas with elevated human-density originally hosted ∼17% of large herbivore biomass and 9% of small herbivore biomass.

## Discussion

### Implications for conservation and ecology

High potential biomass of large herbivores overlaps considerably with densely populated regions and croplands. The implications are multiple. In places where rewilding and restoration are desired, our mechanistic model approach provides a baseline for restoring herbivore populations. However, returning to maximum-attainable biomass levels through rewilding initiatives in these productive areas might only be possible where adequate landscape is available. In situations where reintroduction of large mammals might be problematic, the focus may be shifted towards restoring biomass of other functional groups. In regions struggling with habitat loss and species decline, our estimates offer a carrying-capacity benchmark for assessing biomass losses and prioritizing conservation actions. Particular attention should be dedicated to certain functional groups that have very limited biomass remaining. Given REMAP current spatial and taxonomic resolutions in, these implications are relevant primarily to large-scale patterns; for local and regional conservation applications a finer-scale version of the model will be needed. Furthermore, water dependency in mammals has important implications for their conservation. REMAP could be used in future studies to investigate how changes in the global water cycle brought by climate change might affect herbivore populations (Fuller et al., 2016).

The ecosystemic role of smaller species is often overlooked, even though many are threatened (Dirzo et al., 2014). We provide the first global estimates of small herbivore biomass and show that it comprises a significant, and potentially the largest, component of remaining herbivore biomass. Small species likely play important, and entirely different, functional roles in arid and cold regions (Olofsson et al., 2012; Poe et al., 2019), where water availability is low and where large mammals are declining rapidly (Keesing, 1998). The loss and magnitude of small mammals’ functional biomass is comparable to large animals. Thus, small species should receive more attention in studies of ecosystem functioning and biogeochemical cycles and conservation programs to maintain their functional diversity.

### Mammals in Earth system science

We presented the first global classification of all terrestrial mammalian herbivores (Fig. 1). The HFT system describes the functional diversity of thousands of species using a few key eco-physiological traits. Provided trait data are available, this flexible framework could also be applied to create HFTs for extinct mammals, other endotherms, or ectotherms. Other classifications are possible to create HFTs, but the one we developed facilitates the integration with most IPCC-level global vegetation models. The classification provides a new lens for analyzing the mammalian trait space and biogeographic patterns of functional diversity in relation to environmental factors. REMAP is relatively accurate despite the broad spatial resolution and some limitations in its inputs. Finer-scale inputs would improve biomass estimates in highly heterogeneous landscapes such as transition, deserts, or arid zones. Still, REMAP is highly accurate in regions containing 90% of herbivore biomass (Fig. 2 and S6). Because REMAP includes competition and functional diversity, it better captures trophic interactions and eco-physiological responses to environmental conditions compared to allometric or statistical models. It, therefore, can show the cascading effects of declines in certain functional groups, of reduction of resources, or of climate change through physiological stress. For example, it could be applied to disentangle the effects of climate change on small and large herbivores (Fuller et al., 2016).

The global HFT classification and REMAP lay the foundation for showing how mammals influence biogeochemical cycling, greenhouse gas fluxes, and vegetation dynamics. Our current model does not include all feedbacks between animals and ecosystems, but a foreseeable application would be to fully-couple REMAP with global vegeattion models and include invasive species and livestock, which today comprise the largest herbivore biomass pool (Greenspoon et al., 2023). Further, we envision integration with Earth-system models to demonstrate the influence of mammals in biosphere-atmosphere feedbacks. These will be fundamental steps towards assessing the role of mammalian herbivores and other wild animals in ecosystems and climate.

**Supplementary materials** are available for this paper

## Acknowledgments

We thank J. Chang and M. Clauss for constructive comments and feedback on a previous version of the manuscript, Y. Hui for providing MODIS data, F. Maignan, M. Peaucelle, and N. Viovy for the assistance with ORCHIDEE, P. Stevenson for help with fruit production.

## Data availability

The mammal species classification list with the relevant references is available in the supplementary materials Data S1. The raw species data used for the creation of HTFs, the species density data to calculate herbivore biomass, and REMAP model code are available in a Dryad repository at https://datadryad.org/stash/share/UTASeStv6rt2rF5m1EYMoW8Lzts079VKqiO3gb7piC c. All other data including climatic, environmental, and habitat classification were downloaded directly from their respective sources indicated in the methodology and were used as provided.

## Author Contributions

F.B. obtained the funding. F.B. conceived the project with input from P.C. F.B. develop the methodology with input from all other authors. F.B. performed the analysis and prepared the figures. F.B. led the writing and all other authors contributed with editing and feedback.

