## Supplementary for "Trait-based mechanistic approach highlights global patterns and losses of herbivore biomass functional diversity"

**This PDF file includes:**

Supporting methods
Figs. S1 to S11
Tables S1 to S3

**Other Supplementary Materials for this manuscript include the following:**

Data S1

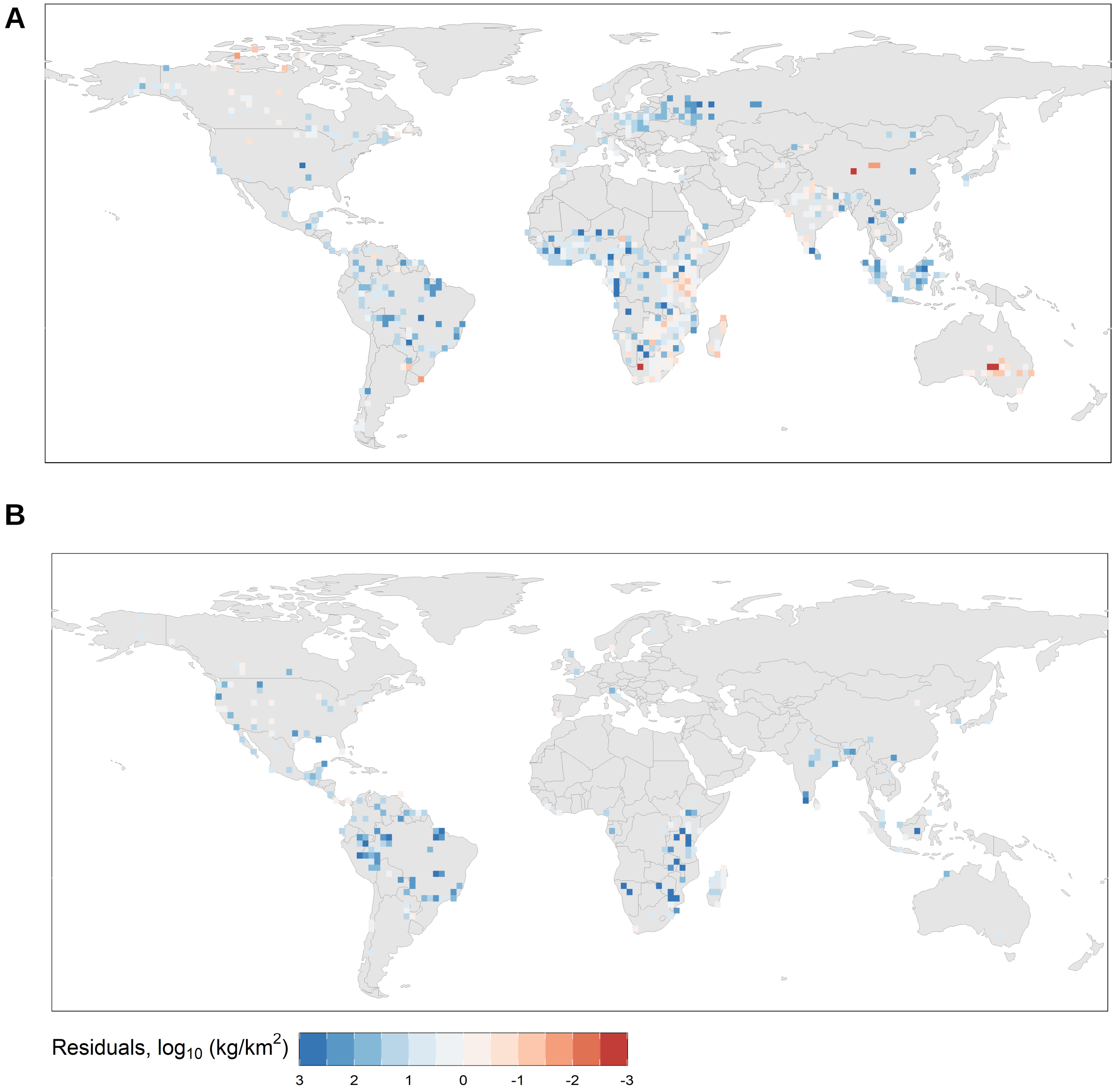


**Fig. S1**. **Distribution of empirical biomass estimates and residuals.** Spatial distribution of biomass residuals calculated from REMAP output minus the empirical-only biomass for large (A) and small (B) herbivores.

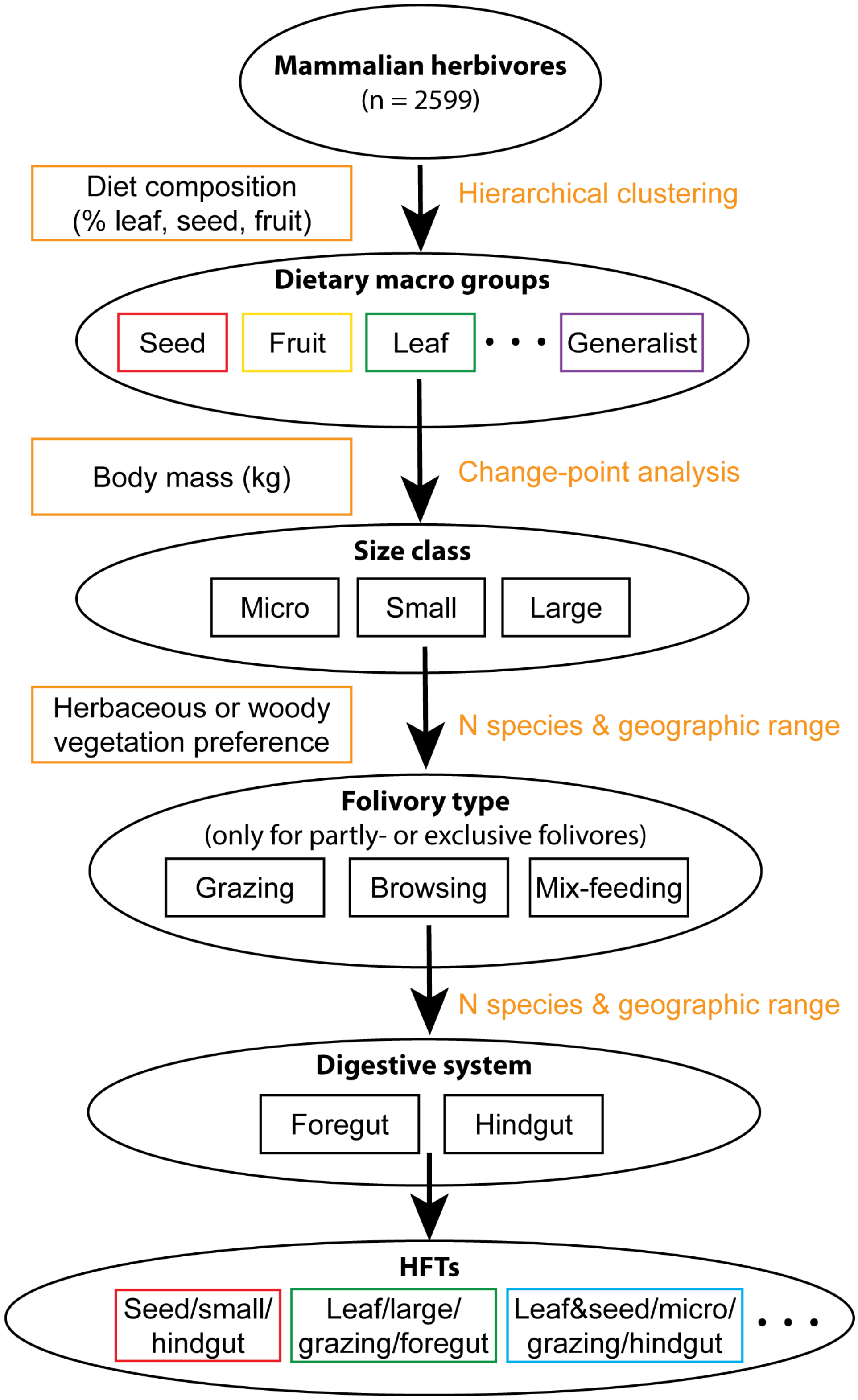


**Fig. S2. Flowchart of creation of HFTs following a nested stepwise procedure.** The orange rectangles indicate the traits used to group species at each step and the orange text indicates the methodology. Not all dietary macro-groups or HFTs are displayed.

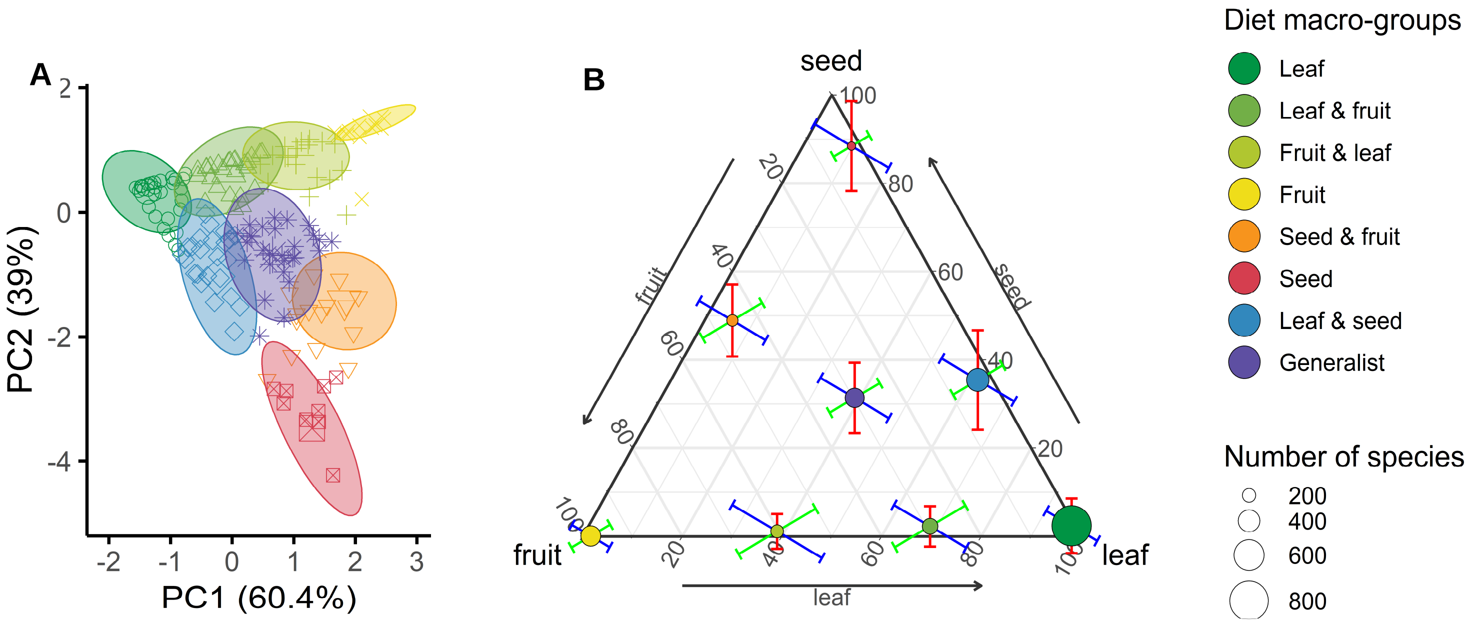


**Fig. S3. Visualization of dietary macro-groups cluster analysis and diet variability.** The order in the name of each macro-group indicates food preference. (**A**) Results of the cluster analysis showing eight distinctive groups. Principle Component Analysis on leaf, fruit, and seed is used to plot the clusters in two dimensions. Relative contribution to PC1: 58% leaf and 41% fruit; PC2: 61% seed and 28% fruit. (**B**) Number of species, diet composition, and variability within macro-groups. Values are expressed in percentages and error bars represent the standard deviation of each plant organ in the diet.

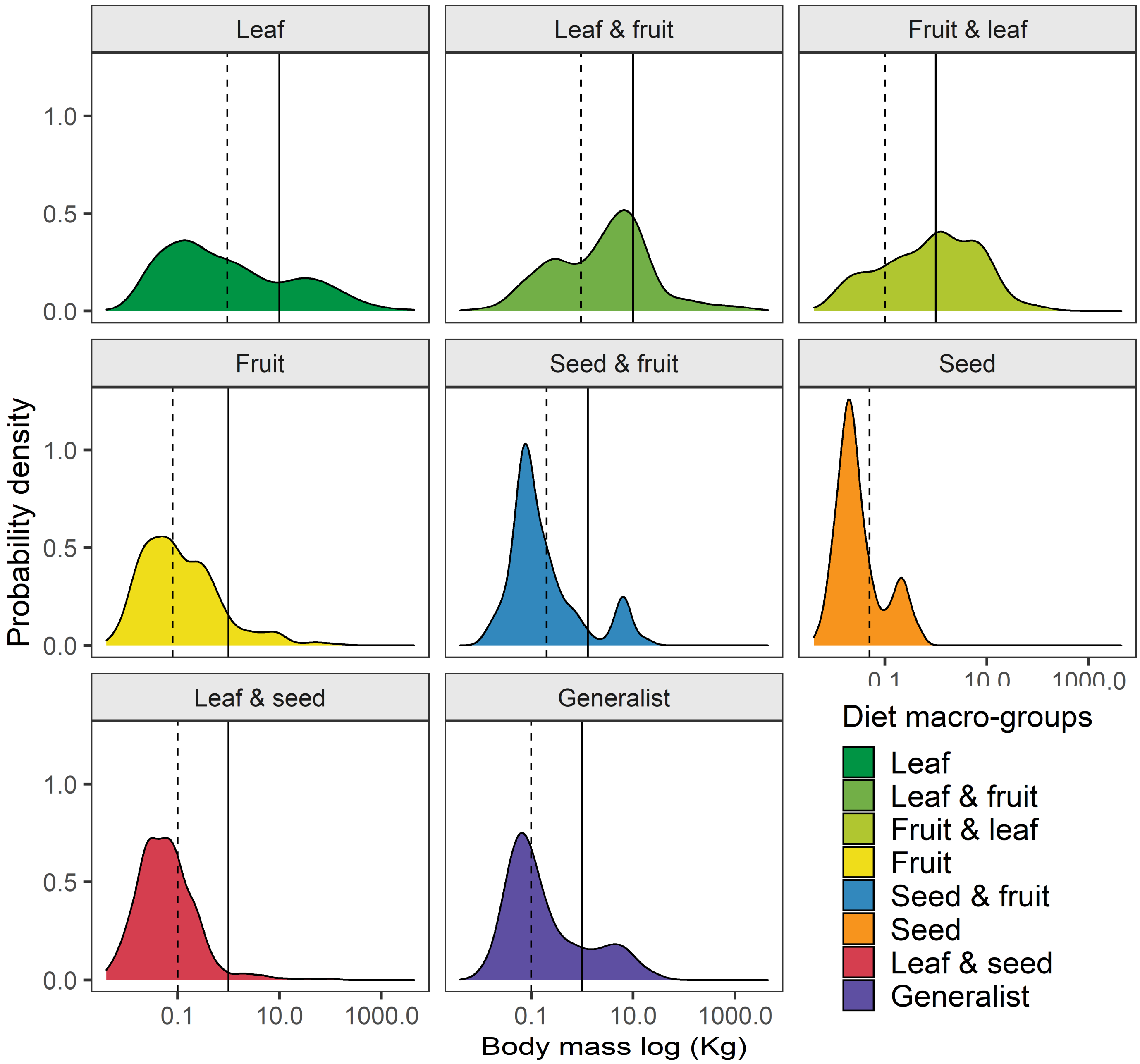


**Fig. S4. Visualization of dietary macro-groups body mass distribution.** Distribution of herbivore body mass across size and functional groups. All herbivores (n=2599) were divided in three size classes. The solid line separates small and large herbivores whereas the dashed line delimits small and micro herbivores.

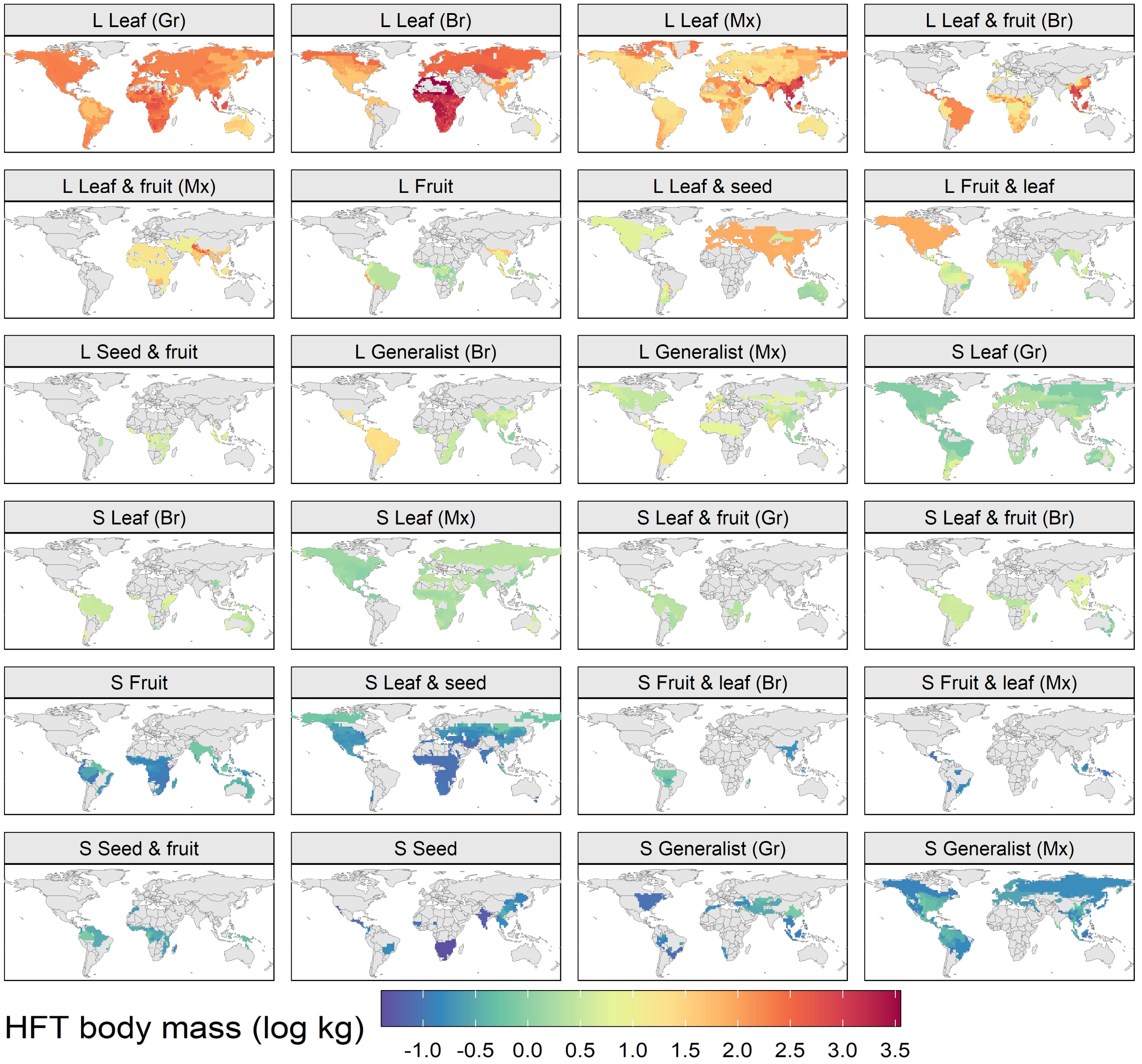


**Fig. S5**. **Body mass of herbivore functional types.** These maps were used to prescribe body mass in the REMAP simulations. The L and S before the HFT name indicate large or small HFTs, respectively. Gr = grazers, Br = browsers, Mx = mixed-feeders

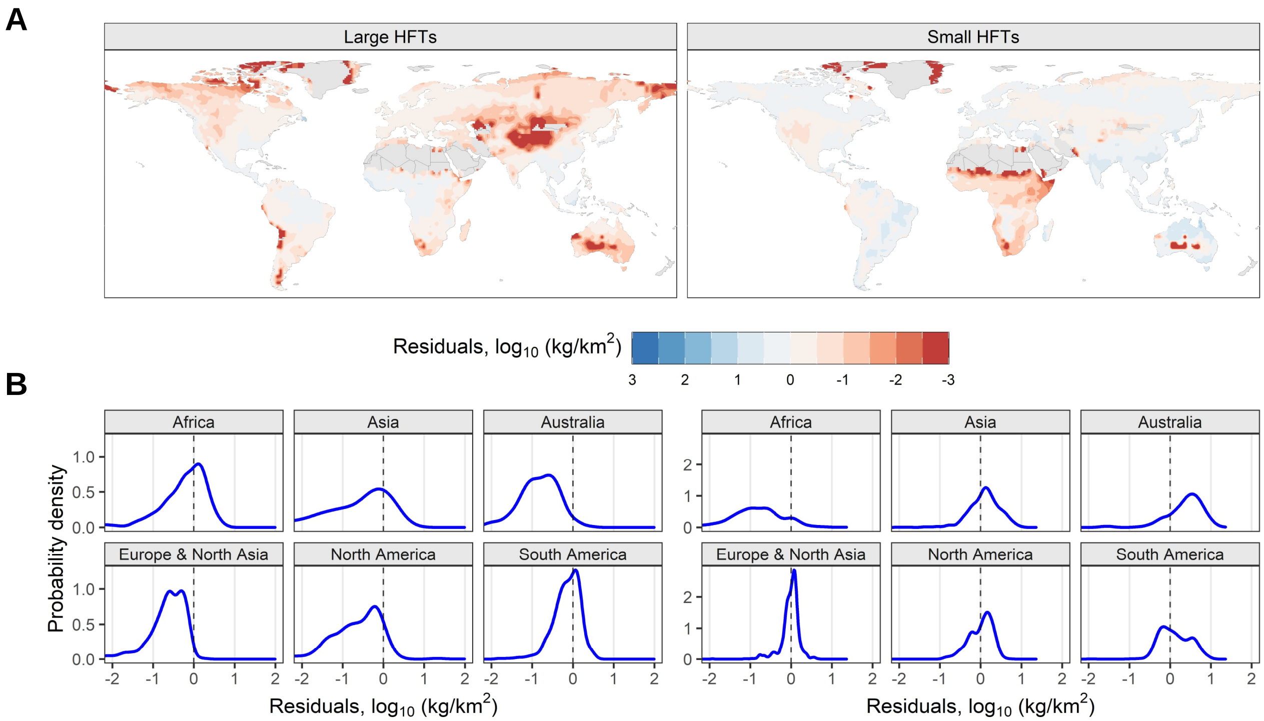


**Fig. S6. Biomass residuals (modeled – empirical). Negative residuals indicate modeled biomass is lower than empirical biomass.** (**A**) Spatial distribution of biomass residuals calculated from REMAP output minus the gap-filled empirical biomass (SI Appendix for details on gap-filling of empirical biomass). (**B**) Probability density of residuals across continents for large HFTs (left) and small HFTs (right)

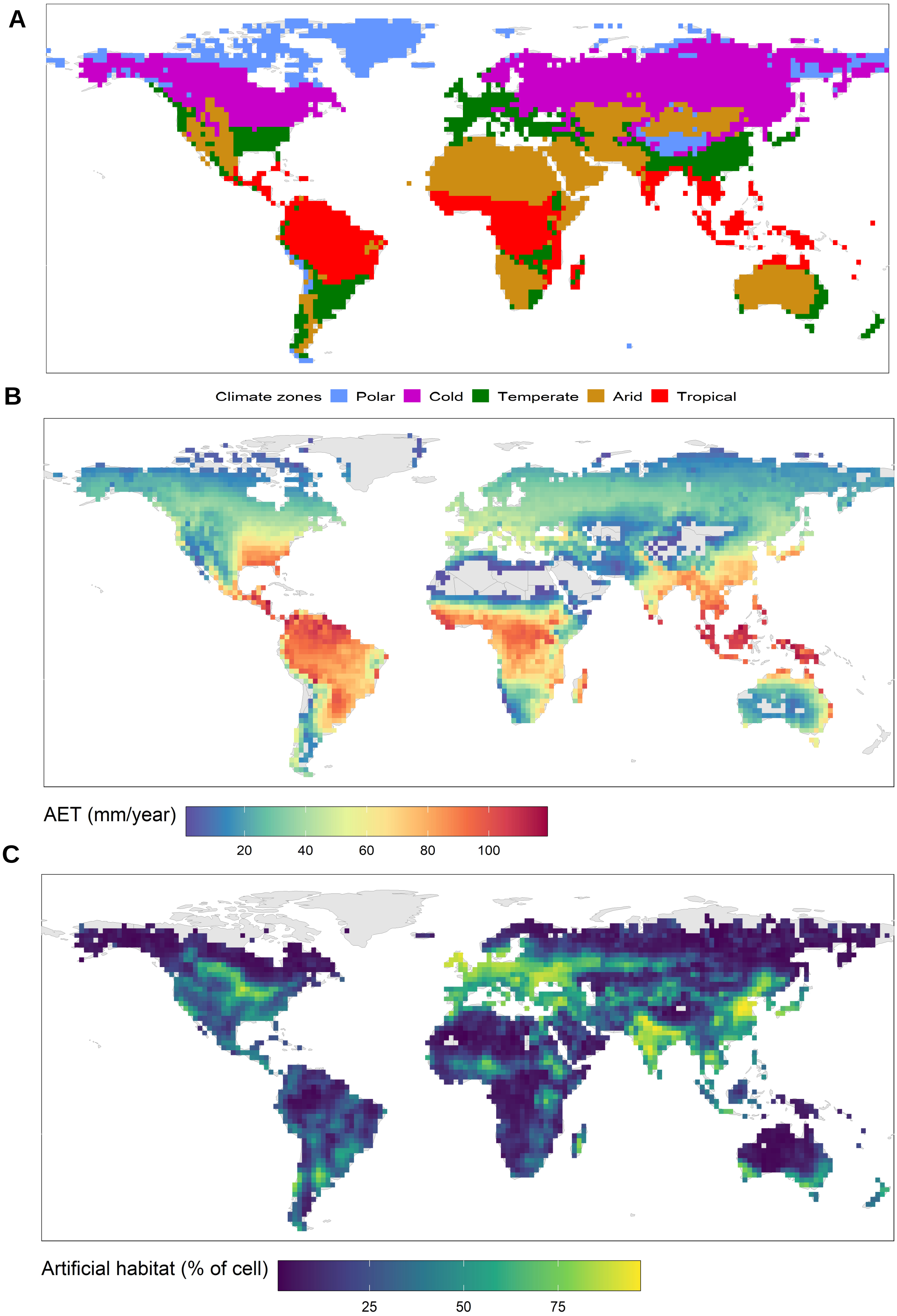


**Fig. S7.** See next page for caption

**Fig. S7. External data used in the analysis.** (**A**) Distribution of climate zones according to the Koppen-Geiger climate classification. (**B**) Plant actual evapotranspiration. The map shows the average of the years 1958-1980[36]. (**C**) Land cover of artificial habitat. Grid cells containing more than 50% of artificial habitat (Arable land, pasture, urban areas, and plantations) according to a global map of terrestrial habitat types (see Methods).

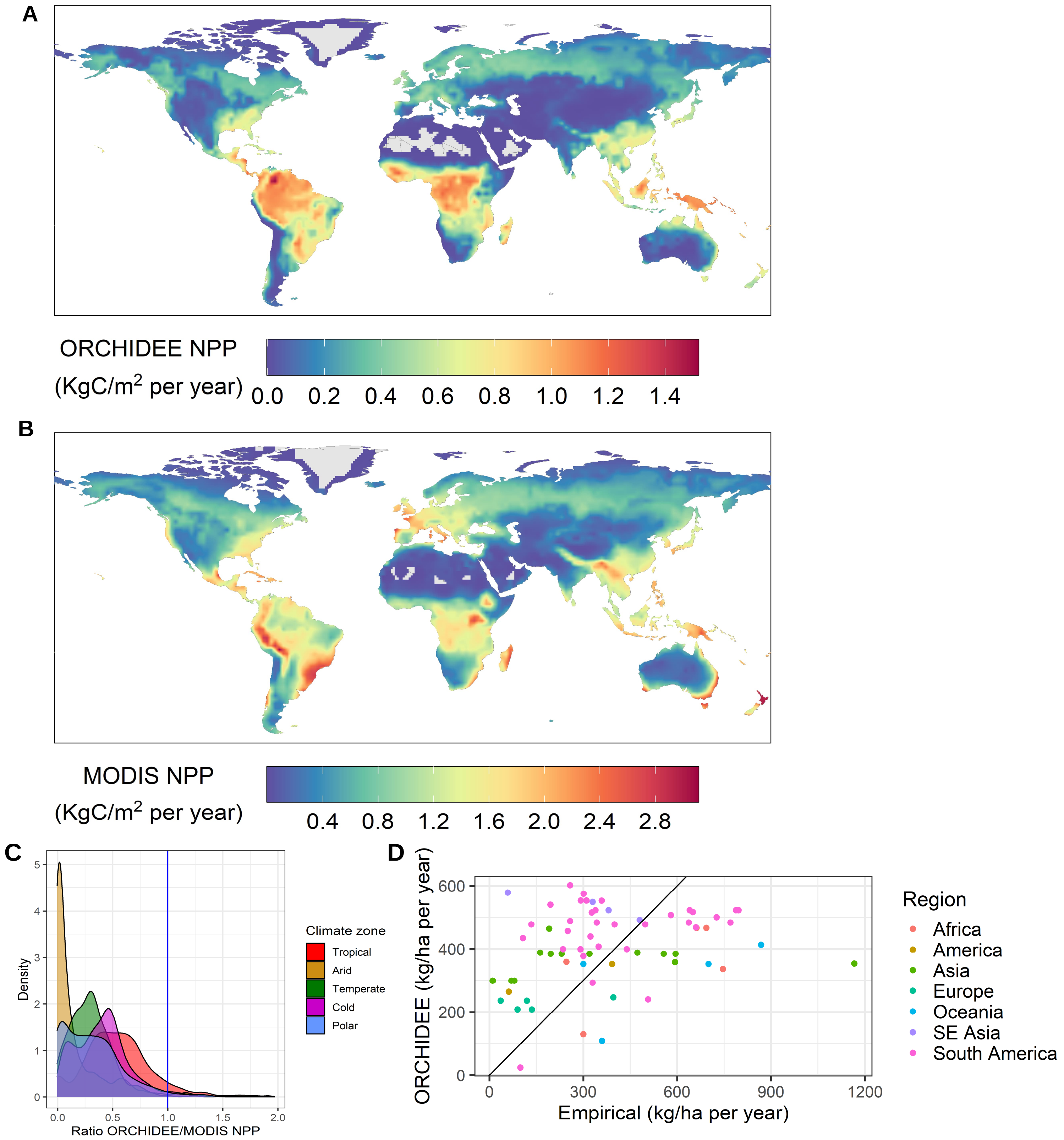


**Fig. S8.** See next page for caption

**Fig. S8. Validation of ORCHIDEE NPP and fruit biomass output.** Comparison of ORCHIDEE and MODIS Net Primary Productivity (NPP): Spatial distribution of (**A**) ORCHIDEE NPP and (**B**) MODIS NPP, note the different range of values between the respective color scales; (**C**) kernel density estimate of the ratio between ORCHIDEE and MODIS NPP across different climate zones. Ideal ratio is 1, ratio < 1 NPP is underestimated, ratio >1 NPP is overestimated by ORCHIDEE. (**D**) Empirical and modeled fruit production. Comparison of fruit fall (dry weight) between the ORCHIDEE model output and empirical measurements

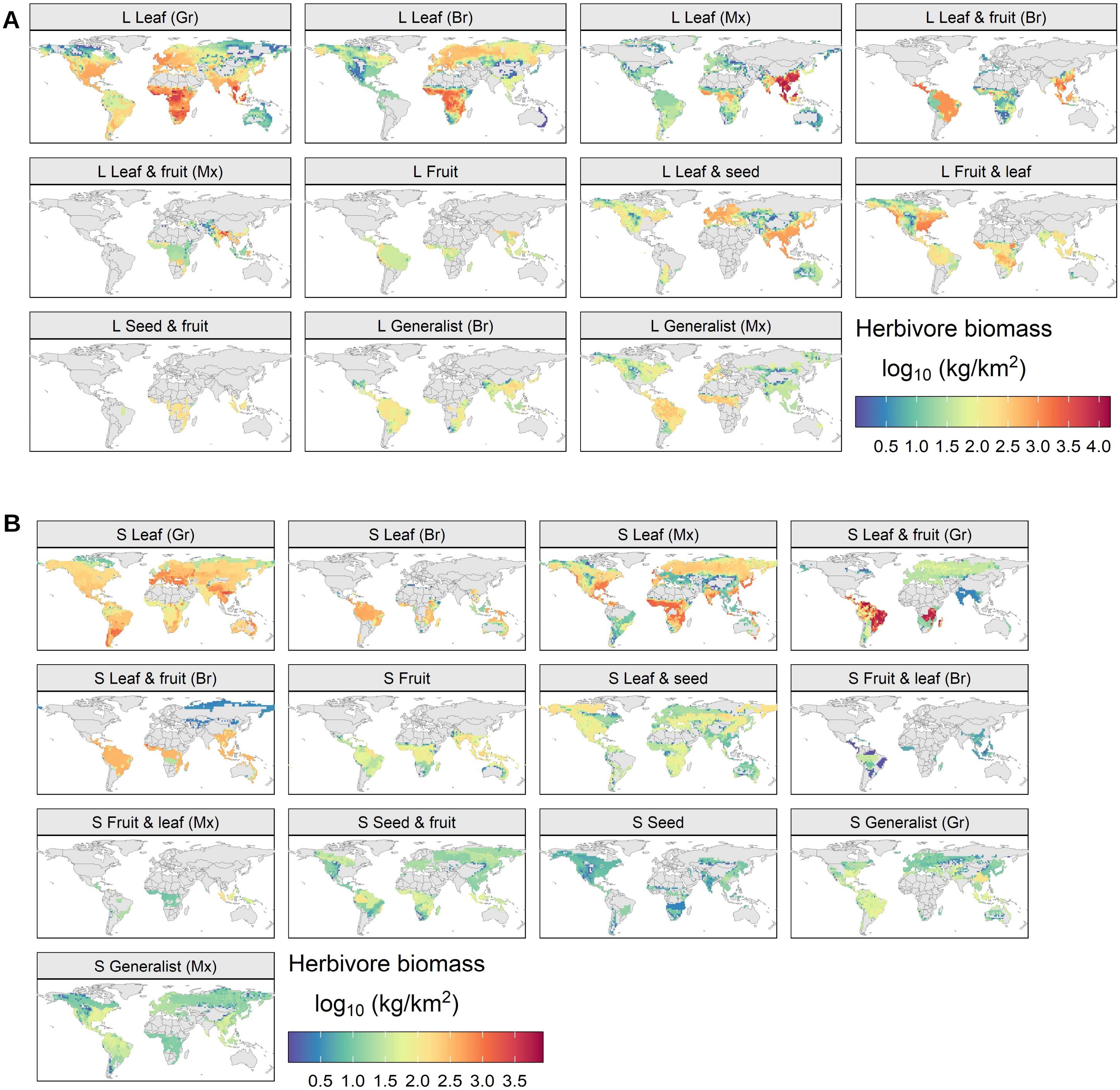


**Fig. S9. Modeled biomass distribution of herbivore functional types under the natural condition scenario**. The L and S before the HFT name indicate large or small HFTs, respectively. Gr = grazers, Br = browsers, Mx = mixed-feeders. (**A**) Large and (**B**) small HFTs.

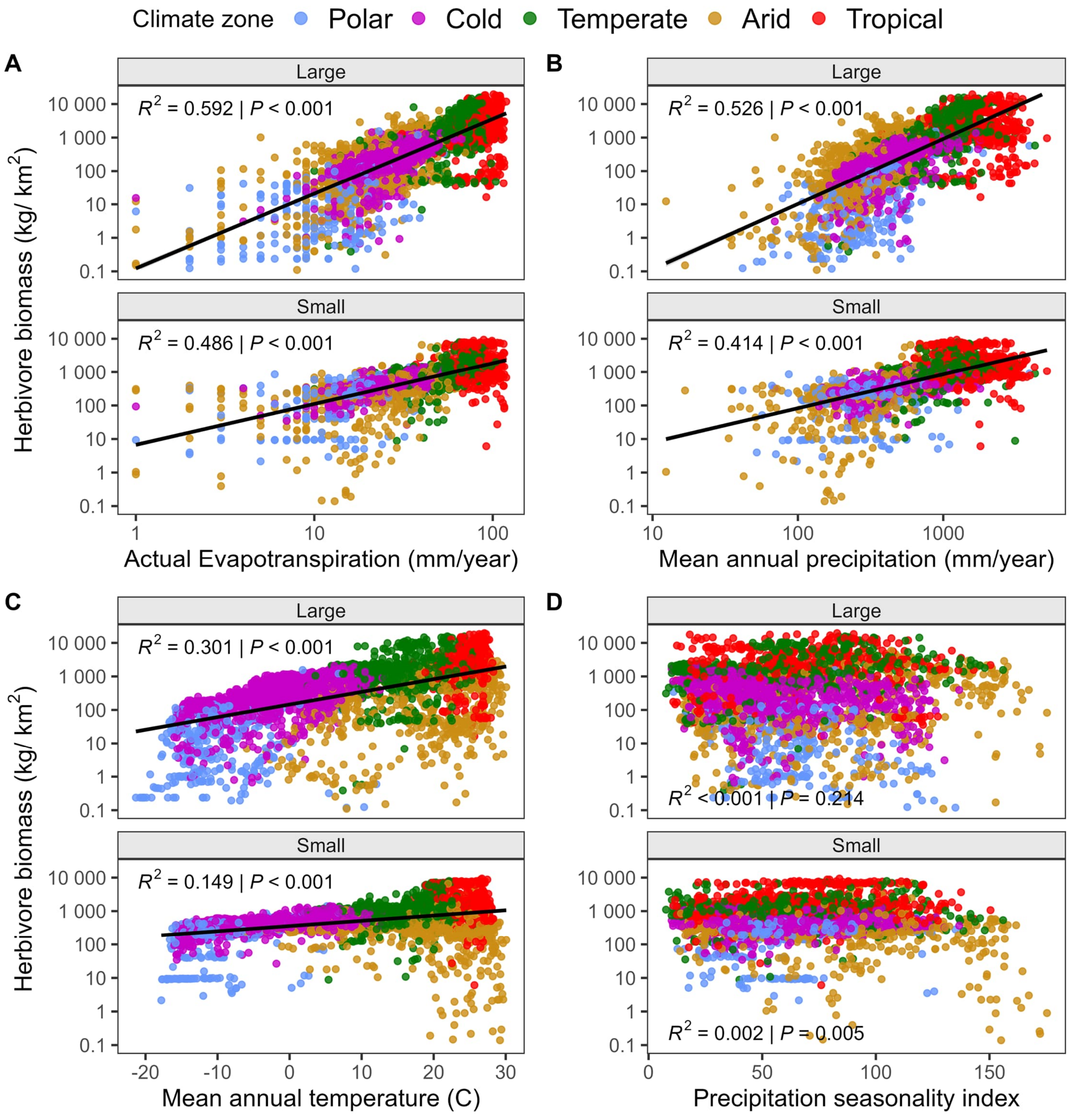


**Fig. S10. Correlations between modeled herbivore biomass under natural conditions and climatic factors.** (**A**) Actual evapotranspiration, (**B**) mean annual precipitation, (**C**) mean annual temperature, and (**D**) precipitation seasonality index, for which regression lines are only shown because of poor fit. Herbivore biomass was log transformed along with actual evapotranspiration and mean annual precipitation.

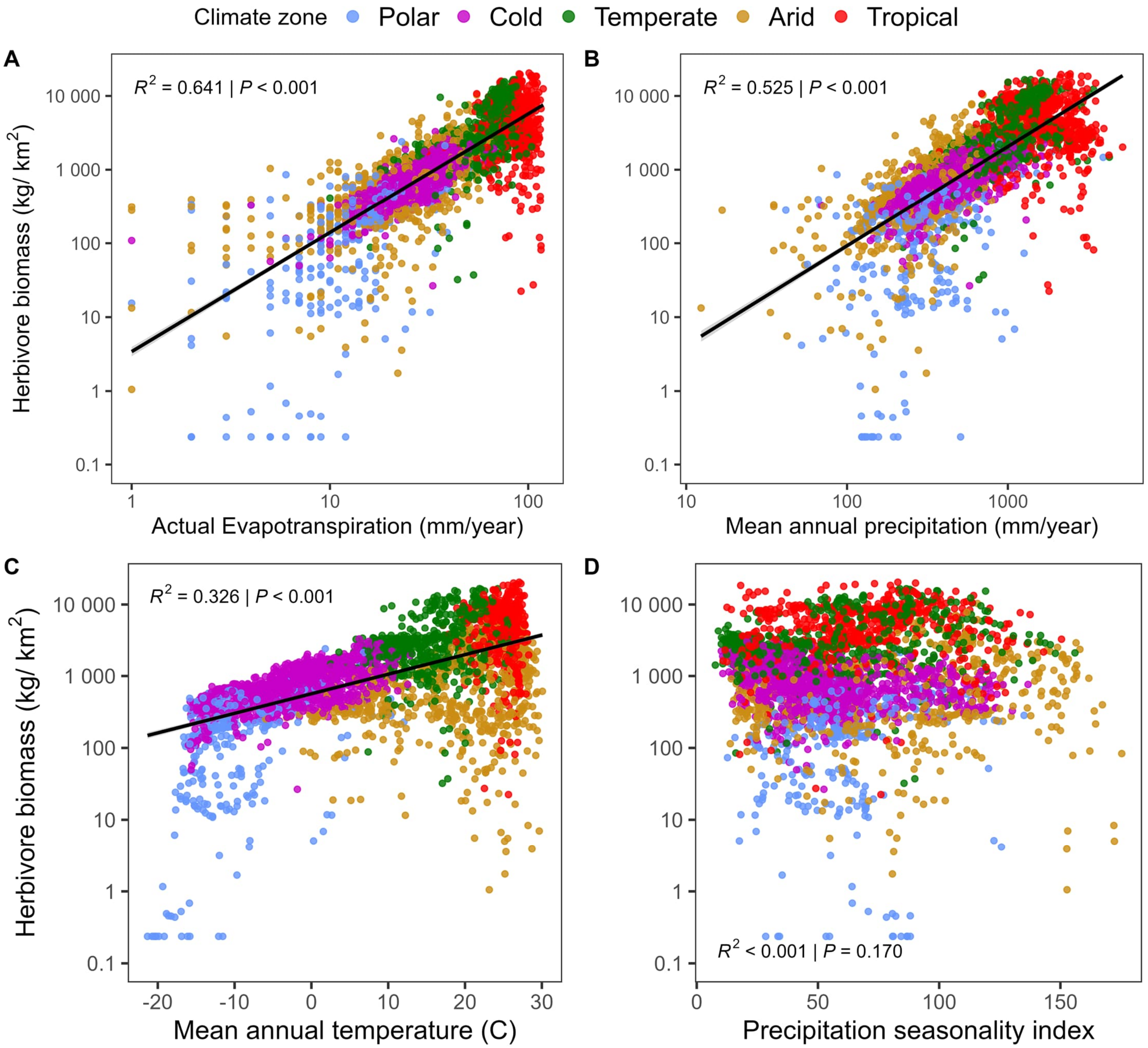


**Fig. S11. Correlations between modeled herbivore biomass under natural conditions and climatic factors.** (**A**) Actual evapotranspiration, (**B**) mean annual precipitation, (**C**) mean annual temperature, and (**D**) precipitation seasonality index, for which regression lines are only shown because of poor fit. Herbivore biomass was log transformed along with actual evapotranspiration and mean annual precipitation.

| HFT diet macro-group | Feeding strategy | Digestive system | | Large HFT total biomass (Kg) | Small HFT total biomass (Kg) | Large HFT mean BM (Kg range) | Small HFT mean BM (Kg range) | N species | |
|---|---|---|---|---|---|---|---|---|---|
| | | L | S | | | | | L | S |
| Leaf | Grazing | F | H | 1.0E+11 | 7.7E+10 | 155 (9-1906) | 1.45 (0.65 - 7.2) | 80 | 58 |
| | Browsing | H | H | 7.0E+10 | 8.4E+09 | 402 (7-3520) | 3.21 (1.16-8) | 31 | 39 |
| | Mixed-feeding | F | H | 7.9E+10 | 3.7E+10 | 61 (7-2222) | 2.1 (0.92-6.8) | 84 | 52 |
| Leaf & fruit | Grazing | - | H | - | 1.3E+10 | - | 2.38 (0.72-7.76) | - | 23 |
| | Browsing | H | H | 1.2E+10 | 1.6E+10 | 111 (7-1231) | 3.73 (0.72-7.76) | 30 | 93 |
| | Mixed-feeding | F | - | 8.4E+09 | - | 20 (8-360) | - | 29 | - |
| Fruit | - | H | H | 2.5E+09 | 1.8E+10 | 7 (1-98) | 0.29 (0.06-0.75) | 34 | 139 |
| Leaf & seed | Grazing | - | H | - | 2.6E+10 | - | 0.27 (0.08-0.67) | - | 112 |
| | Mixed-feeding | H | - | 1.1E+10 | - | 42 (1-81) | - | 13 | - |
| Fruit & leaf | Browsing | H | H | 1.3E+10 | 9.5E+08 | 57 (1-545) | 0.44 (0.17-0.79) | 88 | 28 |
| | Mixed-feeding | - | H | - | 1.2E+09 | - | 0.17 (0.08-0.79) | - | 33 |
| Seed & fruit | - | H | H | 1.7E+09 | 1.1E+10 | 5 (2-15) | 0.39 (0.16-1.04) | 18 | 29 |
| Seed | - | - | H | - | 2.4E+09 | - | 0.11 (0.04-0.34) | - | 20 |
| Generalist (Leaf & fruit & seed) | Grazing | - | H | - | 7.9E+09 | - | 0.22 (0.08-0.69) | - | 39 |
| | Browsing | H | - | 4.7E+09 | - | 9 (1-25) | - | 37 | - |
| | Mixed-feeding | H | H | 3.6E+09 | 1.6E+10 | 5 (1-18) | 0.22 (0.08-0.79) | 31 | 61 |

**Table S1. Eco-Physiological traits.** Traits, biomass, and body mass of HFTs implemented and modeled in REMAP. Digestive system: F = foregut, H = hindgut. The subheadings L and S indicate respectively large and small HFTs. Thicker lines divide the eight dietary macro-groups.

| HFT diet macro-group | Feeding strategy | Digestive system | | Daily intake parameters (i, j , k in eq. 1) | | BM-metabolic rate slope ($k_3$ in eq. 4) | | Density-dependent mortality ($k_d$ in eq. 4) | | Metabolizable energy (MJ kgDM$^{-1}$ eq. 3) | |
|---|---|---|---|---|---|---|---|---|---|---|---|
| | | L | S | L | S | L | S | L | S | L | S |
| Leaf | Grazing | F | H | 0.034 3.57 0.077 | 0.108 3.28 0.08 | 0.73 | 0.686 | 16 | 746 | 11.5 | 11.5 |
| | Browsing | H | H | 0.114 4.013 0.048 | 0.114 4.018 0.048 | 0.734 | 0.613 | 7 | 38347 | 10 | 10 |
| | Mixed-feeding | F | H | 0.057 3.678 0.067 | 0.111 3.65 0.064 | 0.73 | 0.655 | 37 | 4362 | 10.6 | 10.74 |
| Leaf & fruit | Grazing | - | H | - | 0.108 3.28 0.080 | - | 0.707 | - | 16073 | - | 9.82 |
| | Browsing | H | H | 0.098 3.97 0.051 | 0.094 3.969 0.051 | 0.727 | 0.673 | 31 | 4176 | 9 | 9.08 |
| | Mixed-feeding | F | - | 0.034 3.69 0.069 | - | 0.723 | - | 74 | - | 9.6 | - |
| Fruit | - | H | H | 0.114 4.02 0.048 | 0.114 4.02 0.048 | 0.7 | 0.598 | 89 | 735 | 6.6 | 6.6 |
| Leaf & seed | Grazing | - | H | - | 0.108 3.28 0.080 | - | 0.733 | - | 1106 | - | 12.48 |
| | Mixed-feeding | H | - | 0.111 3.65 0.064 | - | 0.704 | - | 73 | - | 12 | - |
| Fruit & leaf | Browsing | H | H | 0.114 4.02 0.048 | 0.114 4.02 0.048 | 0.688 | 0.647 | 117 | 93 | 7.8 | 8.02 |
| | Mixed-feeding | - | H | - | 0.111 3.65 0.064 | - | 0.733 | - | 766 | - | 8.34 |
| Seed & fruit | - | H | H | 0.114 4.018 0.048 | 0.114 4.018 0.048 | 0.62 | 0.73 | 124 | 401 | 11.75 | 11.75 |
| Seed | - | - | H | - | 0.114 4.018 0.048 | - | 0.733 | - | 327 | - | 16.9 |
| Generalist (Leaf & fruit & seed) | Grazing | - | H | - | 0.108 3.284 0.080 | - | 0.733 | - | 619 | - | 11.54 |
| | Browsing | H | - | 0.114 4.02 0.048 | - | 0.691 | - | 114 | - | 10.88 | - |
| | Mixed-feeding | H | H | 0.111 3.65 0.064 | 0.111 3.65 0.064 | 0.718 | 0.733 | 90 | 472 | 11.44 | 11.96 |

**Table S2. REMAP parameters for the herbivore functional types.** Individual parameters for HFTs daily intake, metabolic rate, carrying capacity, and metabolizable energy. The corresponding parameter names and equation are indicated in parentheses.

**A**

| HFT name | Proportion of plant organs in diet (lower bound - upper bound) | | | |
|---|---|---|---|---|
| | **grass leaf** | **tree leaf** | **fruit** | **seed** |
| Leaf grazer | 100 | - | - | - |
| Leaf browser | - | 100 | - | - |
| Leaf Mixed-feeder | 50 (20-80) | 50 (20-80) | - | - |
| Leaf & fruit browser | - | 73 (90) | 27 (10) | - |
| Leaf & fruit mixed-feeder | 36.5 (10-80) | 36.5 (10-80) | 27 (10) | |
| Fruit | - | - | 100 | - |
| Leaf & seed | 35.5 (10-80) | 35.5 (10-80) | - | 29 (10) |
| Fruit & leaf | - | 40 (60) | 60 (40) | |
| Generalist browser | - | 38 (25-46) | 33 (25-46) | 29 (25-42) |
| Generalist mixed-feeder | 19 (7.6-38.4) | 19 (7.6-38.4) | 33 (25-46) | 29 (25-42) |

**B**

| HFT name | Proportion of plant organs in diet (lower bound - upper bound) | | | |
|---|---|---|---|---|
| | **grass leaf** | **tree leaf** | **fruit** | **seed** |
| Leaf grazer | 100 | - | - | - |
| Leaf browser | - | 100 | - | - |
| Leaf mixed-feeder | 50 (20-80) | 50 (20-80) | - | - |
| Leaf & fruit grazer | - | 67 (90) | 33 (10) | - |
| Leaf & fruit browser | - | 72 (90) | 28 (10) | |
| Fruit | - | - | 100 | - |
| Leaf & seed grazer | 64 (90) | - | - | 36 (10) |
| Fruit & leaf browser | - | 42 (40-60) | 58 (40-60) | - |
| Fruit & leaf | 21 (10-50) | 21 (10-50) | 58 (40) | |
| Seed & fruit | - | - | 50 (40-60) | 50 (40-60) |
| Seed | - | - | - | 100 |
| Generalist grazer | 42 (25-47) | - | 30 (25-47) | 28 (25-45) |
| Generalist mixed-feeder | 17.5 (7-38) | 17.5 (7-38) | 27 (25-40) | 38 (25-48) |

**Table S3**. **Diet composition of HFTs.** (**A**) Large and (**B**) small HFTs. Lower and upper bounds were determined based on the diet of all species within each HFT.

**Data S1** in file Supplementary_Table1_HFT_classification.xlsx
Terrestrial mammalian herbivore species with their corresponding traits, HFT, and dietary macro-group. Taxonomy, body mass, and diet source were taken from Phylacine 1.2. Digestive system, feeding type, size class, dietary macro-group, and HFT were determined in this study.
